# Structure of autosynthetic models of balanced cell growth and numerical optimization of their growth rate

**DOI:** 10.1101/2020.09.19.304998

**Authors:** Deniz Sezer, Peter Schubert, Martin J. Lercher

## Abstract

Genome-scale reaction network models are available for many prokaryotic organisms. Yet, to predict the proteome and metabolome of the cell from them, additional information about (i) the nonlinear enzyme kinetics and (ii) the regulation of protein expression by metabolic signals is necessary. Knowledge about the latter could be sidestepped by assuming that expression regulation has evolved to achieve the protein composition that maximizes cellular growth rate. A general mathematical framework for optimizing the growth rate of models comprising an arbitrarily complex metabolic network and a relatively simple protein-synthesis network was recently formulated independently by two research groups [de Groot et al., PLoS Comput. Biol. **16**, e1007559 (2020); Dourado & Lercher, Nature Commun. **11**, 1226 (2020)]. Here, this formalism is further developed with particular focus on carrying out the optimization numerically. To this end, we identify the concentrations of the enzymes as the independent variables of the optimization problem and propose novel multiplicative updates for the iterative calculation of the dependent metabolite concentrations. The reduced gradient method, with analytical derivatives, is employed for the numerical optimization. Additionally, the roles of the dilution of the metabolite concentrations by growth and the commonly invoked constraint on the cell dry mass density are clarified. These developments should lay the basis for the practical optimization of large-scale kinetic models, thus formally connecting the physiological “macrostate” of the cell, characterized by its growth rate, to its “microstate”, described by the cell proteome and metabolome.

**Author summary:** An evolving population of non-interacting, unicellular organisms in a constant environment will maximize its growth rate. By expressing the growth rate as a mathematical function of the cellular composition, it becomes possible to formulate an optimization problem whose solution yields the cell proteome and metabolome at the maximal growth rate. The formulation and solution of such an optimization problem has the potential to elucidate fundamental optimality principles in living cells and to enable the engineering of complex biological systems. Building on previous work, here we address the task of solving this optimization problem numerically. In the process, we elucidate the mathematical role of some common simplifying approximations. This allows us to organize many of the existing formulations of the optimization problem into a hierarchy, whose lower levels are reached by invoking these approximations.

## I. Introduction

Unicellular organisms rely on networks of biochemical reactions to synthesize all their molecular components from a handful of small molecules.^1^ Each reaction in the network has two main attributes: the stoichiometry of the molecules participating in it and the identity of the enzyme that catalyzes the reaction. For many prokaryotes, the enzymes encoded in the genome and their corresponding reactions are currently known at close to a complete genomic coverage.^2,3^ Turning this detailed knowledge of cell metabolism into a predictive tool of cell physiology is a central aim of metabolic engineering and prokaryotic systems biology.

At the core of the mathematical analysis of metabolism lies the observation that, at steady state, the total influx and outflux of every compound in the network should exactly balance each other.^4,5^ Together, these mass-balance equalities of all compounds limit the space of the possible reaction fluxes, and thus constrain the flow of matter from the inputs to the outputs of the network. These fundamental constraints, which depend only on the stoichiometries of the reactions and on their connectivity in the network, are expressed most elegantly using matrix formalism.^6,7^

While the existing methods of metabolic analysis differ greatly in the way this mathematical core is extended, they fall roughly into two main categories: constraint-based modeling^8^ and kinetic modeling.^9^ The former methods supplement the metabolic reaction network with an objective function (typically a linear combination of the fluxes), whose optimum is intended to correspond to the physiological state of the organism under the modeled growth condition.^4,5^ To describe cell growth, the network is typically augmented with an extra “biomass reaction”, which reflects the stoichiometries of the precursor molecules (as well as the energy and reducing equivalents) employed in the synthesis of the cellular macromolecules.^10,11^ When predicting maximum growth, for example, one aims to maximize the flux through the biomass reaction, subject to the mass-balance constraints and to physiologically realistic bounds on the fluxes of the input reactions (e.g., glucose uptake). This general optimization framework, commonly referred to as flux-balance analysis (FBA),^12^ is widely used to calculate optimal reaction fluxes in large metabolic networks. Being unaware of the necessary expression levels of the enzymes and the intracellular metabolite concentrations, however, the optimization may favor costly metabolic routes that are avoided by the cell.^13^

The cell metabolome and proteome take center stage in the kinetic modeling of metabolism, where the flux of every reaction is written as a mathematical function of the concentrations of the metabolites participating in it and the corresponding enzyme. Although the functional form of this relation, known as the rate law, may be very complex if mechanistic details are fully accounted for, simpler generic expressions appear to be sufficient for the analysis of metabolic reaction networks.^14,15^ Even these, however, contain biochemical parameters (e.g., the turnover number of each enzyme and the Michaelis constants of its substrates and products), whose knowledge is currently limited to a relatively small number of well-characterized enzymes of only a few organisms.^16,17^ To further complicate matters, the conditions of the *in vitro* measurements collected in databases like BRENDA^16^ and SABIO-RK^17^ may differ from those of the intracellular milieu, which in turn is likely to be inhomogeneous across the cellular interior. Consequently, the inference of the catalytic parameters by fitting to dynamic time-course or steady-state metabolic data is an essential step in the construction of kinetic models of metabolism.^9^

The mathematical analysis presented in this paper takes as its starting point the constraint-based modeling framework, including an objective function. Differently from FBA and related methods, however, the reaction fluxes are “dressed” with the rate laws of the kinetic modeling, thus enabling the optimization of the objective function directly with respect to the metabolite and enzyme concentrations. Despite the difficulties of obtaining good-quality kinetic parameters mentioned above, the formalism assumes that the rate laws of all reactions in the network, including the numerical values of all necessary rate-law parameters, are known.

In principle, the fluxes calculated with FBA-like methods can be related to the molecular concentrations in a post-processing step by inverting the mathematical relationships of the rate laws.^18^ The solution to this inverse problem, however, is not unique, as different combinations of metabolite and enzyme concentrations could lead to identical fluxes.^18^ An additional criterion is therefore needed to select among the various possibilities. In their systematic study, Noor et al. minimized the total enzyme demand (i.e., the total enzyme mass density) of the modeled metabolic network at fixed reaction fluxes.^18^ As the enzyme concentrations are readily expressed in terms of the fixed reaction fluxes and the metabolites through the rate laws, the minimization problem was formulated in the space of the metabolite concentrations only.^18^ (Note that by fixing the fluxes one implicitly fixes the rate of biomass production, hence the growth rate.)

A similar minimization of the density with respect to the reactants, keeping the reaction flux fixed, was used to analyze metabolic reactions in isolation.^19^ Differently from ref. 18, in this case the objective function received contributions from the densities of both the enzyme and its substrates. The minimization predicted a non-linear relationship between the optimal enzyme and substrate concentrations, which was independent of the magnitude of the constrained flux. This relationship was demonstrated to hold, without any fitting parameter, for the enzyme-substrate pairs of reactions that are the dominant sink of their substrates, and which thus could legitimately be treated as being approximately isolated.^19^

The minimization of the total enzyme mass density at a fixed reaction flux (i.e., fixed growth rate) had also been studied for general metabolic networks in refs. 20 and 21, where the optimal solutions were shown to be elementary flux modes.^6,22,23^ In the context of cell growth, however, the biological motivation for minimizing a mass density at fixed growth rate may not be immediately clear. An objective function with a more direct biological interpretation is the cell growth rate, as it is equivalent to evolutionary fitness for non-interacting unicellular organisms in a constant environment.^24,25^

Previously, Molenaar et al. had argued that many of the observed metabolic choices of prokaryotes could be rationalized as strategies for achieving growth-rate maximization.^26^ As the authors had nicely explained, however, maximizing the growth rate is only possible in the case of models that account for both metabolism and macromolecular synthesis, even if in a coarse-grained manner.^26^ Any model of only part of the complete cellular design, no matter how detailed, does not provide access to the growth rate.^26^ In the light of this insight, many researchers have employed a minimal complete-cell model, with one reaction for metabolism and one reaction for protein synthesis, whose growth rate is maximized analytically.^27–30^ In each case, however, the analytical optimization was carried out in a way that did not generalize to large-scale metabolic models.

Unifying the separate mathematical ingredients outlined above, Dourado and Lercher recently formulated a growth-rate optimization problem for metabolic models of arbitrary size.^31^ First, the enzyme-cost perspective of Noor et al. was integrated with the constraint-based optimization framework by explicitly modeling the enzymes as products of a dedicated biochemical reaction, akin to the biomass reaction of FBA. A new enzyme, the “ribosome”, was assigned to this reaction. As a result, the growth rate of the model becomes a function of the kinetic rate of the ribosome and the amounts of the enzymes that need to be synthesized. Second, the roles of the mass density and the growth rate, as an objective function and a constraint, were exchanged compared to the treatments of refs. 18 and 19. Nevertheless, like there, the problem of optimizing the growth rate was formulated in the space of the metabolite concentrations. This led to closed form expressions for the growth rate, the enzyme concentrations, and the reaction fluxes as functions of the concentrations of the metabolites.^31^ These were further utilized to relate the marginal fitness benefits of all intracellular concentrations, which were shown to be identical in the optimal balanced growth state.^31^

A similar mathematical synthesis was reported simultaneously by de Groot and co-workers,^32^ who modeled the production of proteins in a more general way. Whereas in ref. 31 all enzymes were produced through a single reaction, a separate protein-synthesis reaction was introduced for each enzyme in ref. 32. This seemingly minor generalization prevents the derivation of some of the closed-form expressions of Dourado and Lercher, as we show later below. The main focus in ref. 32 was on extending the geometric picture of a feasible flux cone,^6,7^ which plays a central role in the linear program of FBA, to the nonlinear optimization problem at hand. In particular, “elementary growth modes” were defined as the edges of the nonlinear cone, in analogy to the elementary flux modes in the linear problem.

In the present paper, we work with the more general model of de Groot et al.^32^ With the benefit of hindsight, we show that the principal difference between the formalisms of refs. 31 and 32 is the choice of the independent variables used to parametrize the feasible states of the optimization. As already mentioned, these were the metabolite concentrations in ref. 31. In ref. 32, on the other hand, the feasible states were visualized as being in the space of the ribosome fractions allocated to the synthesis of the separate proteins. We find that while this choice more closely reflects the degrees of freedom accessible to the cell’s regulatory system, it unnecessarily complicates the mathematical expressions needed for the actual numerical optimization, including the analytical derivatives of the growth rate. An equally general alternative is to treat the concentrations of the proteins as independent variables. In our opinion, this choice leads to simpler mathematical expressions that are easier to interpret intuitively. The ensuing optimization framework, developed below, can be viewed as a generalization of resource balance analysis (RBA), in which the metabolite concentrations are absent altogether.^33,34^ In our treatment the metabolite concentrations influence the saturation of the enzymes, and thus appear as the dependent variables of the optimization. This generalization leads to non-convex optimization problems in cases involving the choice among alternative active metabolic subnetworks.^35^

The rest of the paper is organized as follows. A model of a cell in balanced growth, comprising a set of mass-balance equalities, is formulated in Sec. II A. While this is essentially the model of ref. 32 (without a density constraint), we make use of matrix notation, thus making the formalism more concise and easier to code using vectorized objects. In Sec. II B, we pose the problem of optimizing the growth rate, and use the mass-balance equalities of the enzymes to eliminate the ribosome allocation fractions. (These variables were retained in ref. 32, where the concentrations of the proteins were eliminated instead.)

The remaining mass-balance equalities of the metabolites constrain the space of feasible concentrations. In Sec. II C 1 we propose novel multiplicative updates to solve these constraints numerically for the concentrations of the metabolites, treating the concentrations of the enzymes as parameters. While parametrizing the feasible states in terms of the protein concentrations is always possible, using the concentrations of the metabolites as parameters becomes an option when the stoichiometric matrix of the model has additional structure (Sec. II C 2). The requirements that enable such metabolite-centered perspective, first developed in ref. 31, are spelled out in Sec. II E, after introducing three minimal models used to illustrate the developed formalism (Sec. II D). In Sec. II F, we highlight the inability of the examined model to keep its protein concentrations finite upon maximization of the growth rate, and discuss the role of the metabolites in this context. The problem of infinite protein concentrations is overcome by either neglecting the dilution of the metabolites completely (Sec. II G) or by assuming that either the dry-mass density or the macromolecular density of the cell is constant (Sec. II H). In all cases, the reduced gradient method is shown to be well suited for the numerical optimization of the growth rate (Sec. II G 1). The final section of the paper (Sec. III) summarizes our findings, charts the formal relationship between the optimization problem we study and many of its variations in the literature, and ends with an outlook.

## II. Methods and results

### A. Formulation of the model

We model all cellular processes using the mathematics of chemical reaction networks.^36,37^ Consequently, even processes that lack chemical steps, like the passive transport of sugar molecules across the cell membrane, for example, are referred to as “chemical reactions”. We assume that the reactions are catalyzed by enzymes, whose kinetics are specified by appropriate rate laws.^14^ Our treatment neglects stochastic fluctuations that are both extrinsic and intrinsic to the cell,^38^ where the former refers to the differences in the catalytic parameters of the enzymes from one cell to another, and the latter to the fluctuations in the numbers of the chemical species about their averages for constant values of the catalytic parameters. Additionally, we assume that all reactions proceed homogeneously in time and space, thus ignoring complications related to the cell cycle^39,40^ and cell structure.

A single growing cell increases its volume and mass for some time and subsequently divides. In the present context, the issue of individual cell division is avoided by viewing the extensive variables in the formalism (e.g., the amounts of the compounds and the volume in which the reactions take place) as referring not to a single cell but to a cell culture.^41^ Nonetheless, if all cells in the culture are identical, albeit at different ages since their last division, the intensive variables (e.g., densities and concentrations) are also properties of a single (average) cell. For simplicity, we will speak about such a single cell, even when discussing extensive variables.

#### 1. The cell as a network of chemical reactions

The amounts of the chemical species, collected in the vector ***n*** (mol), change in time because of the chemical reactions as follows:

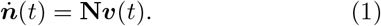

Here, the dot above ***n*** indicates differentiation with respect to time; the vector ***υ*** (mol s^−1^) contains the velocities of the reactions; and the matrix **N** contains the stoichiometric coefficients of the molecules participating in the reactions. (Lowercase and uppercase bold letters are used for vectors and matrices, respectively.)

The mathematical properties of the stoichiometric matrix **N** are of central importance in the structural analysis of the kinetic system (1).^42^ In particular, the left null space of **N** corresponds to quantities that are conserved by the simultaneous kinetics of all reactions, whereas its right null space corresponds to combinations of reaction fluxes that leave the molecular amounts unchanged. While the null spaces of **N** play a fundamental role in the flux-balance analysis (FBA) of metabolic networks,^6,7^ they correspond to properties of the system (1) that are independent of its kinetics. The calculation of the growth rate, however, requires additional, kinetic information.

#### 2. Cell reactions are catalyzed by enzymes

As such, (1) does not constitute a closed dynamical system. To achieve closure, the reaction velocities should be related to the kinetic variables ***n***, whose time derivatives appear on the left-hand side.

Before specifying these relationships, we distinguish between two types of molecules—catalysts (i.e., facilitators of the reactions) and non-catalysts.^31,32^ We will refer to the former as “enzymes” and the latter as “metabolites”. While motivated by the biochemical usage of these terms, here we use them more broadly. In particular, our enzymes will include the ribosome and transporter proteins, without necessarily implying active transport. Similarly, our metabolites will include the substrates and the products of all reactions, as long as the latter are not already classified as enzymes. With this understanding, we partition the vector of moles as

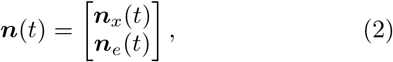

where the subscripts *x* and *e* refer to, respectively, the metabolites and the enzymes. With *N*_*x*_ and *N*_*e*_ denoting the numbers of these two types of molecules, ***n*** has *N*_*x*_ + *N*_*e*_ elements.

Like refs. 31 and 32, we assume that all modeled reactions are catalyzed by enzymes, with the enzymes supposed to be sufficiently dilute to appear linearly in the reaction velocities. Using matrix notation, we write this linear relationship as

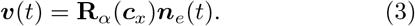

Here, the matrix **R**_*α*_(s^−1^) contains the molar rates of the reactions (i.e., rate per mole of enzyme), which are given by appropriate enzyme rate laws.^14^ These are functions of the concentrations of the metabolites, ***c***_*x*_, and the enzyme kinetic parameters [not indicated in (3)]. Since the concentrations of the chemical species are

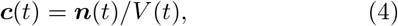

the cell volume, *V* (l, liter), becomes relevant. Although not made explicit in (3), the variables ***c***_*x*_ also depend on the time. (The meaning of the subscript of **R**_*α*_ will become apparent in the next subsection.)

Combining (1) and (3), thus eliminating the reaction velocities, we get

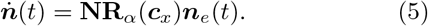

The temporal changes of all chemical species (on the left-hand side of this equality) are driven by the enzymes on the right-hand side. Since the enzymes appear on both sides, the dynamical system (5) is self-replicating^26^ or self-fabricating.^32^ Here, we refer to it as “autosynthetic”, as similar models have been studied before.^43^

Equation (5), together with the definition of the concentrations in (4), is still not a closed dynamical system, since the rate of change of the volume remains unspecified. For full closure, it is necessary to have an additional kinetic equation of the form

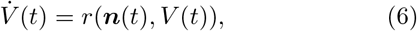

where the rate *r* is, potentially, a function of all kinetic variables. Together, (5) and (6) constitute a closed dynamical system. All questions about its kinetics can be addressed by integrating these differential equations in time.

#### 3. Metabolism and protein synthesis

As in refs. 31 and 32, we will treat all enzymes as proteins and will take all proteins to be enzymes. We lump transcription and translation in an effective protein-synthesis reaction catalyzed by the enzyme “ribosome”. Following ref. 32, we introduce *N*_*e*_ reactions of protein synthesis—one for each enzyme. With the ribosome kept aside, the remaining *N*_*m*_ = *N*_*e*_ − 1 enzymes catalyze the metabolic reactions. For simplicity, we take the metabolic reactions to be in a one-to-one correspondence with the metabolic enzymes.

In line with these assumptions, we partition the vectors in (3) as

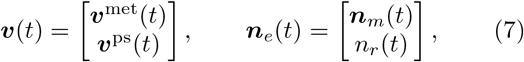

where ***υ***^met^ and ***υ***^ps^ are the rates of, respectively, the metabolic and protein-synthesis reactions, and ***n***_*m*_ and *n*_*r*_ are the amounts of, respectively, the metabolic enzymes and the ribosomes.

While all protein-synthesis reactions are catalyzed by the ribosome, not all ribosomes are simultaneously engaged in the synthesis of the proteins of one particular type. We denote by *α*_*ϵ*_ the fraction of the total ribosome pool allocated to the synthesis of enzymes of type *ϵ*, and collect these fractions in the vector ***α***. For simplicity, we assume that there are no idle ribosomes, hence the sum of the *N*_*e*_ ribosome allocation fractions equals one. This normalization condition is most conveniently written using vector notation as

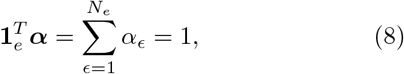

where **1**_*e*_ is a vector composed of *N*_*e*_ ones, and the superscript *T* denotes the transpose of a column vector to a row vector.

With the partitioning of the vectors in (2) and (7), the matrices **N** and **R**_*α*_ partition as follows:

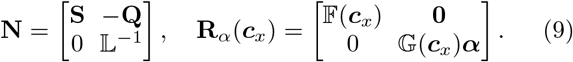

(Blackboard bold style is used for diagonal matrices.)

On the right-hand side of (9), the diagonal matrix 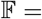 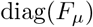, *μ* = 1, … , *N*_*m*_, is composed of the molar rates of the metabolic enzymes, *F*_*μ*_(***c***_*x*_). Similarly, the diagonal matrix 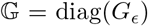, *ϵ* = 1, … , *N*_*e*_, contains the molar rates of the ribosome when working on the synthesis of each of the protein types, *G*_*ϵ*_(***c***_*x*_). As 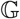 multiplies the vector ***α***, the right half of **R**_*α*_ in (9) is a column vector. The overall dimension of **R**_*α*_ is (*N*_*m*_ + *N*_*e*_) × (*N*_*m*_ + 1).

On the left-hand side of (9), the *N*_*x*_ × *N*_*m*_-matrix **S** is the stoichiometric matrix of the metabolic reactions. The overall normalization of each column of **N** depends on the definition of the reaction velocity in the corresponding row of **R**_*α*_. Differently from ref. 32, we take *G*_*ϵ*_to correspond to the average rate of adding one amino acid to the elongating chain of protein *ϵ*, and not to the production of a complete protein by the ribosome. Because of this choice, the lower right corner of **N** contains the inverse of the matrix 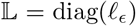, where *ℓ*_*ϵ*_ is the number of monomers (amino acids) making up protein *ϵ*. Concurrently, the *N*_*x*_ × *N*_*e*_-matrix **Q** contains the stoichiometric coefficients of the metabolites participating in the protein-synthesis reactions, but referring to the addition of one amino acid. For the amino acids of the model, the corresponding elements in a given column of **Q** are equal to the fractions of these amino acids in the respective protein. The overall dimension of **N** is (*N*_*x*_ + *N*_*e*_) × (*N*_*m*_ + *N*_*e*_).

Using (9), we write the metabolic and protein-synthesis parts of the kinetic equation (5) as

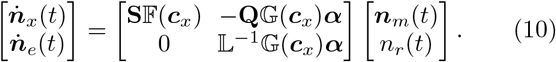

In this dynamical model, the metabolite concentrations, ***c***_*x*_, which appear as arguments of the rate laws in 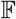 and 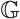, are functions of time, but the ribosome allocation fractions, ***α***, are time-independent parameters.

#### 4. Balanced cell growth

In what sense does the system of differential equations (10) lead to growth? The last scalar equality in (10) is

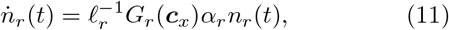

where *ℓ*_*r*_, *G*_*r*_ and *α*_*r*_ refer to the ribosome components of 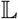, 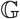 and ***α***, respectively. This equation describes the synthesis of the ribosomes by the ribosomes. As the rate of increase of *n*_*r*_ is proportional to *n*_*r*_, (11) corresponds to exponential growth with instantaneous rate constant

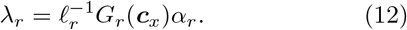

Since more ribosomes will produce more proteins, the numbers of all proteins will increase concurrently. Thus, in our model, cell growth is driven solely by the autocatalytic property of the ribosomes and is independent of the kinetics of the volume.

In reality, the synthesis of proteins and the increase of the cell volume are different, though related, processes that can be decoupled under appropriate experimental conditions.^44^ For a proper understanding of cell size regulation and its role in cell division,^39^ the kinetics of the volume [eq. (6)] should be modeled explicitly.^45^ This, however, is not the purpose of the current paper. Here, we focus on a special type of kinetics, called balanced growth,^46^ in which all molecular constituents and the volume of the cell increase exponentially with constant rate *λ* (s^−1^):

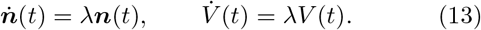

Since one of these equalities sets the growth rate, the remaining *N*_*x*_ + *N*_*e*_ equations act as constraints on the *N*_*x*_ + *N*_*e*_ + 1 kinetic variables of the dynamical system (i.e., ***n***_*x*_, ***n***_*e*_ and *V*). As a result, only one kinetic degree of freedom is left in balanced growth.^47^ This degree of freedom, which can be identified with any linear combination of the kinetic variables, undergoes exponential growth with rate constant *λ*. During the balanced-growth dynamics of the system, the ratios between the time-dependent variables and this reference degree of freedom stay constant. Thus, what remains to be studied is the relationship between *N*_*x*_ + *N*_*e*_ constant ratios and the growth rate *λ*. This is the static (i.e., time-independent) problem that we engage with in the present paper.

From the perspective of chemical intuition, the most convenient choice for a reference kinetic variable is the volume, since the ratios between the other kinetic variables and *V* are equal to the concentrations [eq. (4)]. In balanced growth, all concentrations are independent of time.

The first equation in (13) allows us to eliminate the time derivative on the left-hand side of (5). After dividing the result by the volume, we obtain the following algebraic equation for the concentrations:

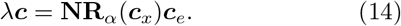

On the right-hand side are the rates with which the concentrations of the compounds would increase due to the activity of the chemical reactions only (i.e., at constant volume). On the left-hand side are the rates with which the concentrations would decrease due to the increase of the cell volume only (i.e., if the reactions were to be switched off). Since the increase of volume dilutes the concentrations of all compounds, we refer to *λ****c*** in (14) as the rate of dilution. In balanced growth, the chemical reactions produce new compounds exactly at the rates that are required to occupy the newly added volume, thereby keeping all concentrations constant.

In the following, we refer to (14) as the balanced-growth equalities of the chemical species. These algebraic equations, whose metabolic and protein parts are [*cf.* (10)]

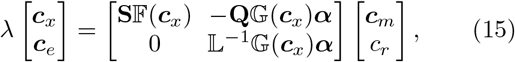

constitute the model of a cell in balanced growth that is studied in the rest of the paper.

### B. The optimization problem

In this section, we consider the problem of maximizing the growth rate of the cell model (15). The optimization is to be carried out with respect to the variables ***c***_*x*_, ***c***_*e*_ and ***α***, which are subject to the balanced-growth equalities (15) and the normalization condition (8). The kinetic parameters in the enzyme rate laws will be treated as fixed.

Defining the vector-valued function [*cf.* (14)]

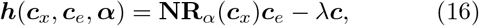

the optimization of the growth rate amounts to the following constrained optimization program:

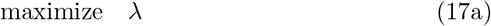

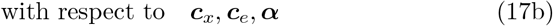

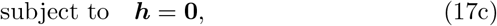

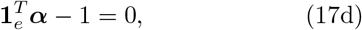

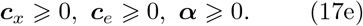

The non-negativity conditions (17e) ensure that the optimization variables (17b) are interpretable as concentrations and ribosome allocation fractions, respectively.

#### 1. Balanced-growth equalities of the proteins

The protein part of the balanced-growth equalities (15) reads

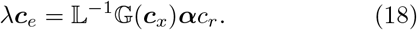

The last of these *N*_*m*_ + 1 scalar equalities recovers the growth rate in (12). In ref. 32, the remaining *N*_*m*_ equalities were used to eliminate the concentrations of the metabolic enzymes from the optimization problem by expressing them in terms of the other optimization variables. (This is shown in Sec. S5 of the SI.)

Here, we will eliminate the ribosome allocation fractions, ***α***. To be able to do so, we assume that the growth rate is strictly positive (*λ* > 0), which immediately implies that some ribosomes (***c***_*r*_ > 0) are allocated to the synthesis of new ribosomes (*α*_*r*_ > 0). Additionally, none of the metabolites that are necessary for the synthesis of the needed proteins should be missing, i.e., if ***c***_*ϵ*_ > 0 for some enzyme *ϵ*, then *G*_*ϵ*_(***c***_*x*_) > 0. To simplify the discussion of the analytical results, below we will take *G*_*ϵ*_(***c***_*x*_) > 0 for all *ϵ*, independently of whether ***c***_*ϵ*_ = 0 or not. This will guarantee that the diagonal rate matrix 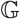 is invertible.

Because the matrices 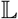 and 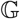 are diagonal, the scalar equalities (18) are decoupled and can be solved for ***α***:

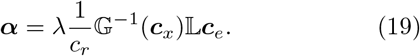

By taking the dot product of (19) with **1**_*e*_, and using the normalization constraint (17d), the growth rate is readily expressed in terms of the concentrations only:

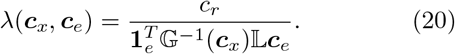

Substituting this growth rate back in (19), we get

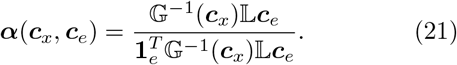

This closed-form expression for the ribosome allocation fractions automatically ensures that ***α*** is properly normalized and *α*_*ϵ*_ ⩾ 0.

#### 2. Balanced-growth equalities of the metabolites

Having used the normalization (17d) and the protein part of (17c) to express the objective function (17a) and the variables ***α*** in terms of the concentrations, we are left only with the metabolic part of (17c) as an equality constraint.

Substituting ***α*** from (19) in the metabolic part of the balanced-growth equalities (15), the latter can be written as

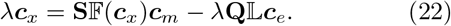

The two additive contributions on the right-hand side of (22) correspond to the rates of production of the metabolites by the metabolic reactions 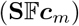 and to their consumption in the reactions of protein synthesis 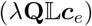. The rate of metabolite dilution is on the left-hand side.

Since the growth rate *λ* is a function of the concentrations [eq. (20)], the *N*_*x*_ scalar equalities (22) are non-linear in both the metabolites and the proteins. Hence, an analytical elimination of *N*_*x*_ optimization variables is not possible in general. How to do this numerically will be discussed in the next section.

Having eliminated the variables ***α***, we transform the original optimization problem (17) to the following equivalent program:

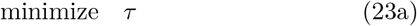

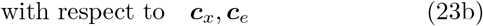

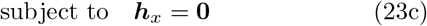

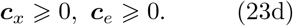

Here, we have defined the objective function (23a), which is to be minimized, as the reciprocal of the growth rate in (20):

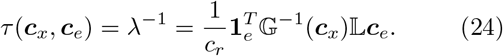

We will refer to *τ* as the “growth time” of the cell. Additionally, in (23c) we defined the vector-valued function

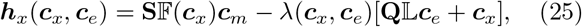

which imposes the metabolic balanced-growth equalities (22) as constraints to the optimization.

Although the non-negativity conditions (23d) appear as separate inequality constraints, in numerical work it is possible to impose them automatically by variable transformation.^48^ The bounds *c*_*i*_ ≥ 0 can be handled by working with the variables *y*_*i*_, defined as 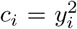, instead of *c*_*i*_. In the case of strict inequalities, *c*_*i*_ > 0, it is possible to use *y*_*i*_ = ln *c*_*i*_. Indeed, working with the logarithms of the metabolite concentrations in the numerical optimization of metabolic networks is a common practice.^18^

The optimization problem (23) is in the space of the *N*_*x*_ + *N*_*e*_ concentration variables (23b). Since these are constrained by the *N*_*x*_ scalar equalities (23c), the feasible space of the problem has *N*_*e*_ degrees of freedom, which can be used to parametrize all balanced-growth states of the model.

### C. States of balanced growth

In this section we discuss how to numerically solve the *N*_*x*_ equality constraints (23c) for some of the variables in terms of the others. First, we treat the *N*_*e*_ enzyme concentrations as the independent variables and determine the *N*_*x*_ metabolite concentrations from them. Then, we consider the opposite case, namely treating the metabolite concentrations as the independent variables and solving for the protein concentrations.^31^

#### 1. Protein concentrations as parameters

To impose the constraint (23c), with the protein concentrations treated as parameters, we view ***h***_*x*_ in (25) as a function of the metabolite concentrations only and look for its root, ***c***_*x*_. This is a multidimensional root-finding problem, which can be solved numerically.

##### a. Additive update rule

The best-characterized multidimensional root-finding method is Newton-Raphson.^49^ In our case, it consists of updating the metabolite concentrations iteratively, until convergence, as

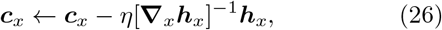

where the *N*_*x*_ × *N*_*x*_-matrix **∇**_*x*_**h**_*x*_ is the Jacobian of the function ***h***_*x*_ with respect to the variables ***c***_*x*_, and the “learning rate” *η* adjusts the step size. (The Jacobian is given in the Sec. S1B of the SI.) At every iteration step, ***h***_*x*_ and **∇**_*x*_***h***_*x*_ need to be re-calculated at the new values of ***c***_*x*_, and the Jacobian needs to be inverted.

Being an additive update rule, (26) can lead to negative, and hence meaningless, concentrations during the iterations. As already mentioned, one way of preventing this is to work with the logarithms or square roots of the metabolite concentrations.^48^ Another possibility is to replace the additive update by a multiplicative update, as described next.

##### b. Multiplicative update rule

Multiplicative updates have been employed successfully in optimization problems with non-negative optimization variables. First used in the non-negative factorization of matrices,^50^ they were applied more recently to quadratic programming with non-negativity constraints^51^ and polynomial root finding.^52^ Given the non-negativity conditions (23d), multiplicative updates are highly relevant to our case.

The equality constraint (23c) reflects the balance between inward fluxes that increase the concentrations of the metabolites and outward fluxes that decrease these concentrations. As these fluxes contribute additively to the constraint function (25), the latter is of the form

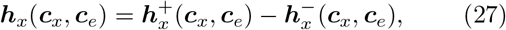

where the elements of the vectors 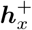 and 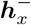 are non-negative for all choices of the concentrations **c**_*x*_ and ***c***_*e*_.

Because of this special form of ***h***_*x*_, its root ***c***_*x*_ can be obtained using the following multiplicative update rule:

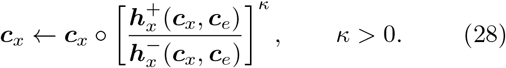

Here, the multiplication (denoted by ○), the division, and the power function are elementwise operations on the vectors. Since the ratios 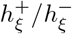 in (28) are non-negative for all *ξ* = 1, … , *N*_*x*_, the metabolite concentrations are guaranteed to remain non-negative throughout the iterations. The parameter *κ* plays a role similar to the learning rate *η*, and can be varied (e.g., around *κ* = 1) to improve the stability (smaller *κ*) and convergence rate (larger *κ*) of the algorithm.

The separation (27) is readily achieved by splitting the matrices **S**, 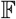, and **Q** in (25) into their positive and negative parts as

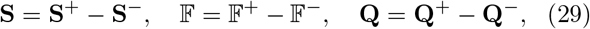

where all elements of **S**^±^, 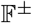, and **Q**^±^ are non-negative. Then, 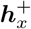 and 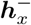 are [*cf.* (25)]

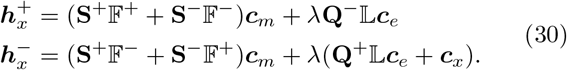

Note that 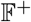 and 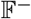 contain the contributions of, respectively, the forward and backward directions of the reversible reactions. Since **S**^+^ has non-zero entries for the products of the forward reactions, and **S**^−^ has non-zero entries for the products of the backward reactions, 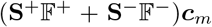 contains the rates of *production* of the metabolites by the metabolic reactions. The last additive contribution to 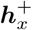 in (30) accounts for the production of the metabolites (e.g., ADP) by the reactions of protein synthesis. Similarly, the vector 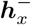 contains the rates of *consumption* (including the dilution, *λ***c**_*x*_) of the metabolites. Thus, according to (28), if 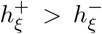 (i.e., the rate of production of metabolite *ξ* is larger than its rate of consumption) the concentration *c*_*ξ*_ should increase. Conversely, if 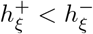 (i.e., the rate of consumption is larger than the rate of production) the concentration *c*_*ξ*_ should decrease. Qualitatively, this mimics the actual response of the reaction network. The update terminates when, for all metabolites, the rates of production are equal to the rates of consumption, which is the state of balanced growth. For the right-hand side of (28) to be finite, the consumption rates, 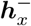, of all present metabolites should be strictly positive. (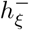 can be equal to zero when *c*_*ξ*_ = 0.)

The physical interpretation of 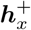 and 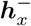 as rates of production and consumption justifies the condition *κ* > 0 in (28). On the basis of the separation (27) alone, we could equally well have decided to multiply ***c***_*x*_ by the reciprocal ratio 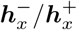 (i.e., *κ* < 0), which is also guaranteed to be non-negative. This latter choice, however, would increase the concentration of a metabolite when its rate of production is smaller than its rate of consumption, resulting in a destabilizing positive feedback.

The multiplicative update (28) has the practical appeal of rendering obsolete the differentiation of the enzyme rate laws and the subsequent inversion of the Jacobian matrix, which were required by the Newton-Raphson update (26).

While multiplicative updates are simple to implement, formally establishing their convergence properties has proven more difficult. Since most such analyses are concerned with applications to non-negative matrix factorization,^53–56^ convergence in other contexts is yet to be clarified.^51,52^

#### 2. Metabolites as parameters

When the stoichiometry matrix of the model is of full column rank, it also becomes possible to select the metabolite concentrations as the independent variables that parametrize the states of balanced growth. As this situation was analyzed thoroughly in ref. 31, it is revisited here only to highlight the extra flexibility gained in the presence of additional structure.

By assuming that the columns of the *N*_*x*_ × *N*_*m*_-matrix **S** are linearly independent (i.e., **S** is of full column rank), we effectively limit the discussion in this case to models whose metabolites are more than, or equal in number to, the metabolic enzymes (*N*_*x*_ ≥ *N*_*m*_). Since the rank of **S** is now equal to *N*_*m*_, **S** lacks a right null space but may have a left null space. The two null spaces bear directly on the task of solving the *N*_*x*_ equality constraints (23c) for the *N*_*m*_ concentrations of the metabolic enzymes, ***c***_*m*_.

##### a. The equality constraints

When **S** lacks a right null space, the *N*_*m*_ × *N*_*x*_-matrix

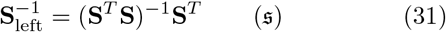

is a left inverse of **S**, i.e., 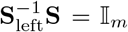, where 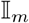 is the *N*_*m*_ × *N*_*m*_ identity matrix. (Equations that apply only to models whose metabolic stoichiometry matrices have a left inverse will be indicated with 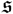.) After both sides of the *N*_*x*_ equality constraints (23c) are multiplied by 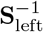 from the left, these are transformed into the *N*_*m*_ equality constraints

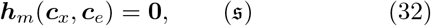

for the vector-valued function

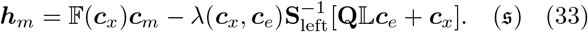

For given metabolite concentrations, ***c***_*x*_, and ribosome concentration, *c*_*r*_, the root of this function, ***c***_*m*_, can be obtained numerically as discussed below.

If there are more metabolites than metabolic enzymes, the left null space of **S** is of dimension *N*_left_ = *N*_*x*_ − *N*_*m*_, and *N*_left_ linearly independent row vectors in this space can be stacked to form the *N*_left_ × *N*_*x*_-matrix **V**_left_ such that **V**_left_**S** = 0. Multiplying both sides of (23c) from the left by **V**_left_, the following additional *N*_left_ equalities are deduced (recall that *λ* > 0):

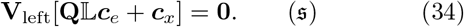

As the rows of **V**_left_ correspond to quantities that are conserved by the kinetics of the metabolic reactions,^7,42^ the conservation realations encoded by (34) relate the concentrations of *N*_left_ (dependent) metabolites to the concentrations of *N*_*m*_ (independent) metabolites.^31^ Consequently, the concentrations of the independent metabolites plus the ribosome concentration provide the full set of *N*_*m*_ + 1 = *N*_*e*_ parameters necessary to parametrize the states of balanced growth.

##### b. Updating the protein concentrations

Viewing ***h***_*m*_ in (33) as a function of the concentrations of the metabolic enzymes, ***c***_*m*_, we can again employ the Newton-Raphson method to find its root. The condition in this case is that the *N*_*m*_ × *N*_*m*_ derivative matrix **∇**_*m*_**h**_*m*_ is invertible. This Jacobian of ***h***_*m*_ with respect to ***c***_*m*_ is worked out in the SI (Sec. S1C). Since **∇**_*m*_**h**_*m*_ contains 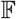 as an additive contribution [*cf.* (33)], it is going to be invertible if the molar rates of all metabolic enzymes are non-zero, i.e., *F*_*μ*_(***c***_*x*_) ≠ 0 for all *μ*. This is the case if all metabolic reactions are active^31^ and none of the reversible reactions is in chemical equilibrium.

In the simple examples that we consider later in the paper, the elements of the vector 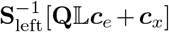 in (33) will be non-negative. Additionally, the diagonal elements of 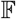 will be non-negative, as our illustrative models will contain only irreversible reactions. Under these simplifying conditions, the multiplicative update that can be used instead of an additive Newton-Raphson update, becomes [*cf.* (33)]

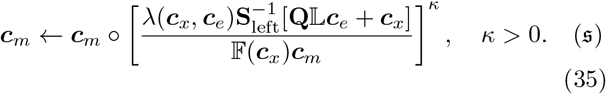

[In general, the matrices 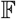, 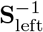 and 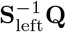 should be written as the difference of two non-negative parts, and the update rule (35) should be modified accordingly.] Since and the division line in (35) denote elementwise operations, for *κ* = 1 the above update simplifies to

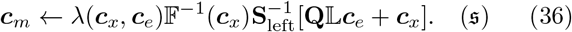

[The same right-hand side can be obtained directly from (32) and (33) after inverting the matrix 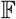 and isolating ***c***_*m*_.]

To justify (35) intuitively, we note that the denominator, *F*_*μ*_*c*_*μ*_, is equal to the flux (per unit volume) catalyzed by metabolic enzyme *μ*. The numerator, on the other hand, contains the rate of dilution of all cell components downstream of this reaction.^31^ For given ***c***_*x*_ and ***c***_*e*_, if the rate of dilution is larger than the reaction flux (i.e., the expression in the square brackets is larger than one), the update increases the concentration of the enzyme, *c*_*μ*_, and hence the flux. Conversely, if the flux of the reaction is larger than the rate of dilution of its downstream compounds, the expression in the square brackets is smaller than one and the enzyme concentration is decreased. The procedure converges when, for every reaction, the dilution of the downstream chemical species is equal to its flux, within a prespecified precision.

### D. Illustrative minimal models

To illustrate the developed theory and provide numerical examples in the subsequent sections of the paper, we introduce the three simple models shown in Fig 1. In these models, *A* and *B* denote the metabolites while *R*, *T*, *U*, and *V* denote the proteins (enzymes). The chemical reactions of the models are given in the second, and their stoichiometric matrices in the third rows of the figure. In accordance with the choice made before, the protein-synthesis reactions reflect the addition of one amino acid to the respective proteins.

**Fig 1.**
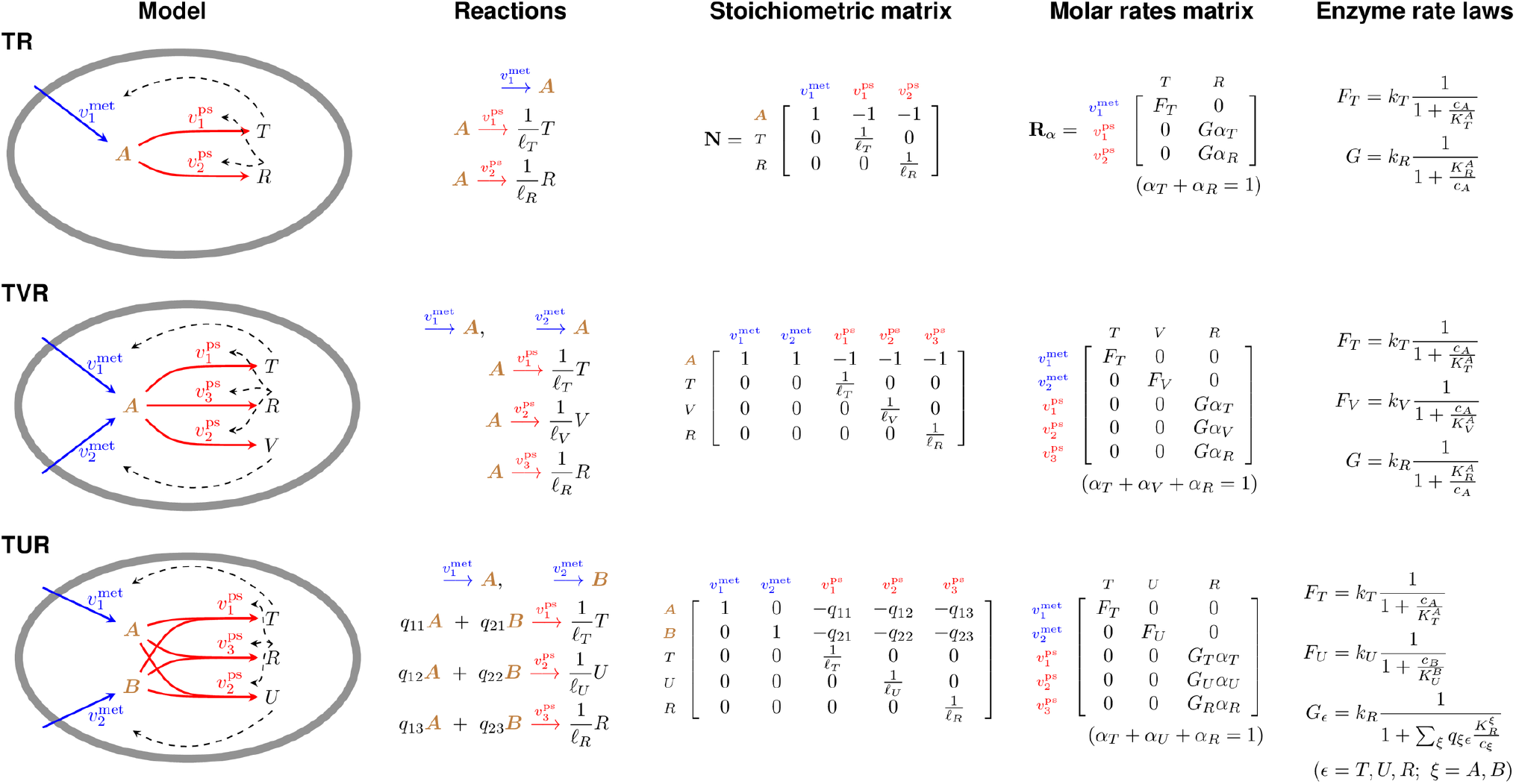
Illustrative minimal autosynthetic cell models. The metabolic reactions are colored blue, the protein-synthesis reactions are red. Dashed arrows connect the enzymes to the reactions that they catalyze. All enzymes are imagined to be proteins.

The models in Fig 1 were selected as the minimal models exhibiting a certain property. The first model, TR, is the minimal possible autosynthetic cell model with separate reactions for metabolism and protein synthesis. Variations of this minimal model have appeared extensively in the literature.^27–31^ The second model in Fig 1, the TVR model, is the minimal autosynthetic cell model with non-square, hence non-invertible, stoichiometric matrix. The proteins in the TR and TVR models are built from a single precursor molecule. As a result, all proteins have identical amino acid composition. The third, TUR model, is the minimal model that allows for proteins with different amino acid compositions. Consequently, it falls outside the class of models analyzed in ref. 31.

The fourth row of Fig 1 contains the molar rate matrices of the models, written here for general rate laws of the metabolic enzymes (*F*_*μ*_) and the ribosome (*G*_*ϵ*_). The particular enzyme rate laws used subsequently in the numerical examples are given in the last row of the figure. They correspond to irreversible reactions but in the case of the metabolic proteins include product inhibition, which accounts for the occupancy of the enzyme active site by the product(s) of the reaction.^57^ In these expressions, *k*_*μ*_ (s^−1^) is the turnover number of metabolic enzyme *μ*, and 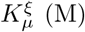 is the Michaelis constant of this enzyme for metabolite *ξ*.

In the left half of Table 1, we have listed possible values for the numbers of amino acids of type *ξ* contained in a protein of type *ϵ*. (These coefficients, 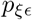, form an *N*_*x*_ × *N*_*e*_-matrix **P**.) The numbers in the table were chosen arbitrarily but considering that the median length of the *E. coli* proteins is approximately 300 amino acids.^58^ Dividing the coefficients 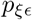 by the total number of amino acids in the corresponding protein (denoted by *ℓ*_*ϵ*_), we obtain the matrix **Q**, which appeared in the upper-right corner of the stoichiometry matrix **N** [eq. (9)]. (Using matrix notation, 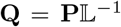) The elements of **Q**, 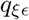, which correspond to the 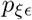 in the left half of Table 1, are given in the right half of the table.

**Table 1.** Assumed amino acid content (**P**) and fractional composition (**Q**) of the proteins in the three minimal models.

|  |  | **P** |  |  | **Q** |  |  |
| --- | --- | --- | --- | --- | --- | --- | --- |
|  |  | *T* | *V/U* | *R* | *T* | *V/U* | *R* |
| TR | *A* | 300 | - | 400 | 1 | - | 1 |
| TVR | *A* | 300 | 350 | 400 | 1 | 1 | 1 |
| TUR | *A* | 200 | 150 | 100 | 2/3 | 3/7 | 1/4 |
|  | *B* | 100 | 200 | 300 | 1/3 | 4/7 | 3/4 |
| | $\ell_\epsilon$ | 300 | 350 | 400 | | | |
| | $60 \times \ell_\epsilon$ | 18000 | 21000 | 24000 | | | |

Since protein synthesis is template-directed polymerization, the rate law of the ribosome is modeled with an equation that is applicable to such a process.^59^ For an abundant template, the molar synthesis rate of a polymer composed of *ℓ*_*ϵ*_ monomers is well approximated by^59^

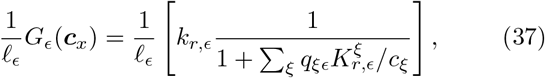

where *k*_*r,ϵ*_ (s^−1^) is the turnover number of the ribosome for attaching *one monomer* to polymer *ϵ*, and 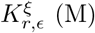 is the Michaelis constant of the ribosome for monomer *ξ* during the synthesis of polymer *ϵ*.

Depending on the level of desired detail, it may be reasonable to assume that the constants *k*_*r,ϵ*_ and 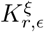 of the ribosome are independent of the type of the synthesized protein, i.e,

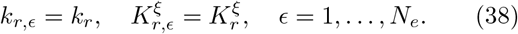

Although not essential for the presented analysis, this assumption was made when writing the ribosome rate law in the last row of Fig 1.

Table 2 contains the values of the enzyme parameters that we use in the numerical examples later in the paper. The catalytic rate constants of the metabolic enzymes *T*, *U* and *V* were chosen based on the observation that the transport rate for sugar transporters saturated with external substrate is approximately 100 s^−1^.^58^ The rate constant of the ribosomes *R* was chosen to be the largest value of the translation rate interval for *E. coli* (10-20 aa s^−1^).^58^ Although otherwise arbitrary, the Michaelis constants were intended to be on the same order of magnitude as the concentrations of the metabolites in *E. coli*.^60^ The *K*_*M*_ values for the ribosomal substrates *A* and *B* were set to, respectively, 10 mM and 5 mM. The *K*_*M*_ values of the metabolic proteins, which reflect the occupancy of the transporters by the intracellular substrates, were chosen to be in the range of tens of mM.

**Table 2.** Turnover numbers (*k*_*ϵ*_) and Michaelis constants 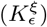 of the enzymes in the illustrative models.

| | $T$ | $R$ | $T$ | $V$ | $R$ | $T$ | $U$ | $R$ |
| --- | --- | --- | --- | --- | --- | --- | --- | --- |
| $k_\epsilon$ ($\text{s}^{-1}$) | 100 | 20 | 100 | 50 | 20 | 100 | 70 | 20 |
| $K_\epsilon^A$ (mM) | 100 | 10 | 100 | 75 | 10 | 100 | - | 10 |
| $K_\epsilon^B$ (mM) | - | - | - | - | - | - | 70 | 5 |

In our models, the cell consists of only two or three different types of proteins. In reality, the total amount of ribosome necessary to incorporate 20 amino acids per second to an extending polypeptide chain should be more than 400 amino acids. Similarly, although *T*, *U* and *V* were imagined to be transporters, the effective enzyme amounts involved in supplying amino acids to the ribosome should be more than 300 or 350 amino acids. With these considerations in mind, we increased the sizes of all enzymes uniformly by a factor of 60. The resulting protein sizes are given in the last row of Table 1. As this rescaling brought the calculated growth rates of the models close to the experimental values for *E. coli* (≤ 2 h^−1^), these larger values will be used as *ℓ*_*ϵ*_ in the numerical examples of the following sections.

### E. Uniformly composed proteins

Even after the assumption that the catalytic properties of the ribosome do not depend on the identity of the synthesized protein [eq. (38)], the molar rates *G*_*ϵ*_ in (37) still depend on the protein type *ϵ* through the stoichiometric coefficients 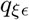. In many coarse-grained models the proteins are either built from only one building block (e.g., the TR and TVR models of Fig 1) or from several building blocks combined in fixed ratios (e.g., the TUR model with 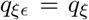 for all *ϵ*). As a result, all proteins in such models have the same amino acid composition, with possible differences in the total number and order of the amino acids.

In this section, we consider such models with uniformly composed proteins to illustrate how the analysis simplifies when the matrix **Q** has additional structure. Although not stated there explicitly, the analysis of ref. 31 applies to such models with structured **Q**.

#### 1. Separable protein-composition matrix

The condition 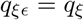 for all *ϵ* has two related implications. First, the composition matrix **Q** has *N*_*e*_ identical columns and is of the separable form

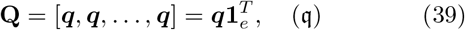

where the vector ***q*** denotes one of the columns. (Equations that hold only for models with uniformly composed proteins will be indicated with 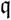.) Second, the molar rates of protein synthesis are independent of the protein type:

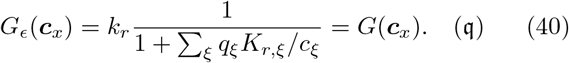

As a result, 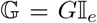, where 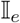 is the *N*_*e*_ × *N*_*e*_ identity matrix.

This special form of 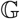 immediately simplifies many of the above equations. For example, the relationship between the concentrations of the protein types and the fractions of the ribosomes allocated to their synthesis [eq. (21)] becomes independent of the kinetic properties of the ribosome:

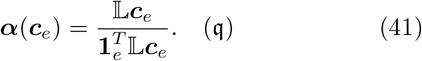

As a result, the ribosome allocation fractions are basically a reweighted (and normalized) version of the protein concentrations.

The vector 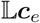 in the numerator of (41) reports the concentrations of the enzymes in units of amino acids. The sum of all these concentrations in the denominator of (41) is the total concentration of amino acid units incorporated in the proteins:

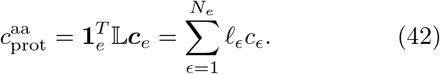

Thus, when all proteins have uniform amino acid composition, the fraction of ribosomes dedicated to each protein (*α*_*ϵ*_) is equal to the fraction of amino acids this protein contains out of the total number of amino acids in the proteome 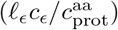.

Another expression that simplifies for 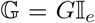 is the growth rate [eq. (20)]:

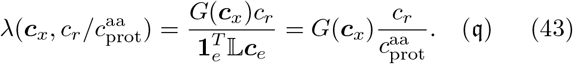

When all proteins have uniform composition, they appear in the growth rate only as the ratio between the number of ribosomes and the number of amino acids in the entire proteome 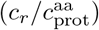.

Other expressions simplify as a result of the separable form of **Q** in (39). Since in this case 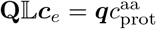, the constraint function ***h***_*x*_ [eq. (25)] becomes

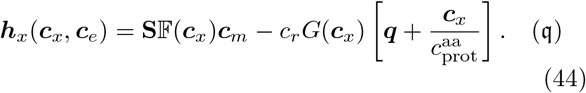

Because of the property (39), we will refer to models with uniformly composed proteins as models with separable **Q**. We will refer to models with general, non-uniform composition of their proteins as models with non-separable **Q**. These two possibilities correspond to the columns of Table 3. The rows of the table distinguish whether the matrix **S** has a left inverse. In this way, the autosynthetic cell models of the type considered in this paper are classified according to the properties of their matrices **S** and **Q**. As an example, the illustrative models from Fig 1 have been assigned to their respective classes in Table 3.

**Table 3.** Classes of autosynthetic cell models according to the specified properties of their matrices **S** and **Q**. Each illustrative model from Fig 1 is assigned to its corresponding class (in parenthesis).

| | | $\mathbf{Q}$ | |
| --- | --- | --- | --- |
|  |  | separable (q) | non-separable |
| $\mathbf{S}$ | left inverse (s) | $\mathbf{s}\mathbf{q}$ (TR) | $\mathbf{s}\cdot$ (TUR) |
| | no left inverse | $\cdot\mathbf{q}$ (TVR) | $\cdot\cdot$ |

#### 2. Invertible metabolic stoichiometry matrix

From Sec. II C 2 we recall that the left inverse of **S** allows for expressing the concentrations of the metabolic enzymes in terms of the concentrations of the metabolites and the ribosome. In the case of models with uniformly composed proteins, the root ***c***_*m*_ of (44) is

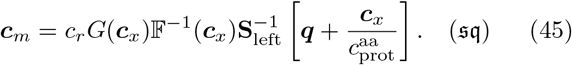

[This expression corresponds to eq. (5) of ref. 31.] If there are more metabolites than metabolic enzymes (i.e., **S** has more rows than columns), the metabolite concentrations should additionally satisfy the *N*_left_ conservation equalities [*cf.* (34)]

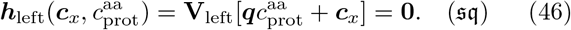

(Since there are no linearly dependent metabolites in the three simple models of Fig 1, an additional example is provided in Sec. S4A to illustrate this situation.)

Using (45), the concentrations of the metabolic enzymes can be eliminated from the objective function (23a). To this end, we rewrite the reciprocal of the growth rate in (43) as

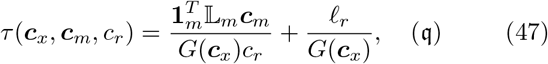

where 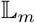 and *ℓ*_*r*_ are the components of 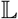 that correspond to, respectively, the metabolic enzymes and the ribosomes. Substituting (45) into (47), we get

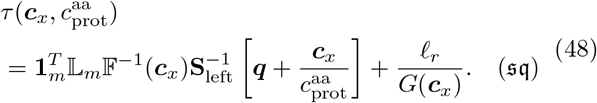

[This result corresponds to eq. (6) in ref. 31.] Importantly, the objective function does not depend on any of the individual protein concentrations but only on the total protein concentration expressed in units of amino acids, 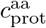.

Because the remaining constraint (46) is a function of the same variables as the objective function (48), in this case the optimization problem (23) simplifies to

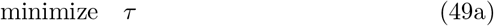

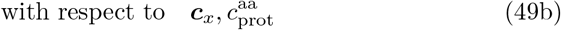

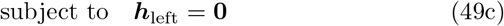

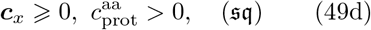

where the proteins are present only in bulk as 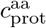. [Up to the assumption of constant dry mass density, which we discuss in Sec. II H 2, the problem (49) is essentially the optimization that was formulated in ref. 31.] This optimization is in the space of *N*_*x*_ + 1 concentration variables that are subject to *N*_left_ equality constraints.

For a square matrix **S**, the equality constraint (49c) drops out completely and the optimization of the growth rate is unconstrained (assuming one works with the logarithms or the square roots of the concentrations).

#### 3. The TR model

Let us illustrate the analysis in the case of uniformly composed proteins by applying it to the TR model. Since there is only one metabolite, the constraint ***h***_*x*_ is a scalar function. With *c*_*T*_, *c*_*R*_ and *c*_*A*_ denoting the concentrations of the two proteins (*T* and *R*) and the metabolite (*A*), we have

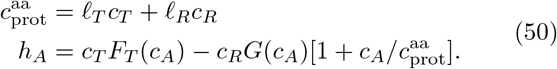

We determined the balanced-growth states of the TR model for *c*_*T*_ and *c*_*R*_ in the range from 1 *μ*M to 300 *μ*M, intended to roughly correspond to measured values in *E. coli*.^61^ (The *k*_cat_ and *K*_*M*_ parameters were as given in Table 2.) Figure 2a shows the *c*_*A*_ values obtained with the Newton-Raphson algorithm. Identical results were obtained using the multiplicative update (28) with *κ* = 1, which for the TR model reads

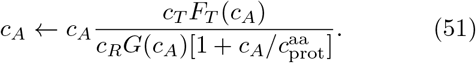

**Fig 2.**
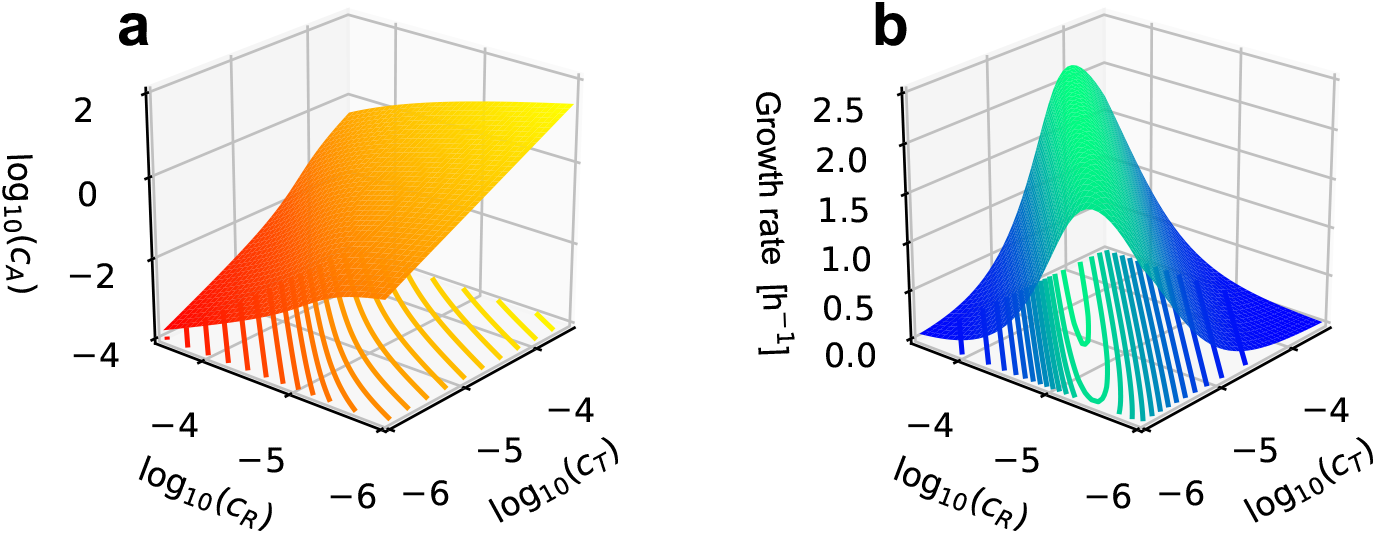
States of balanced growth (**a**) and the corresponding growth rate (**b**) of the TR model. The base-10 logarithms are of the molar concentrations of the ribosome (*c*_*R*_), transporter (*c*_*T*_), and metabolite (*c*_*A*_). The contour lines of constant elevation are shown on the *x*-*y* planes of the plots. The enzyme parameters were fixed at their values in Table 2.

The metabolite concentration in Fig 2a changes by five orders of magnitude in the examined range, while the protein concentrations vary by only 2.5 orders of magnitude. At a fixed ribosome concentration, *c*_*A*_ increases with *c*_*T*_. This makes sense since more metabolite will accumulate inside the cell if the amount of the transporter is increased. In contrast, at a fixed transporter concentration, *c*_*A*_ decreases for increasing *c*_*R*_, as more metabolite is utilized when there is more ribosome.

The growth rate (43) is plotted in Fig 2b. Unlike the metabolite concentration, it is non-monotonic and exhibits a clear maximum. In the examined range of protein concentrations, the maximum appears to be reached at the *c*_*R*_ = 10^−3.5^ M boundary, where *c*_*T*_ ≳ 10^−4^ M.

Interestingly, the contours of constant *λ*, which are drawn on the *c*_*T*_-*c*_*R*_ plane in Fig 2b, run almost parallel to the *c*_*T*_ = *c*_*R*_ diagonal. Since the two horizontal axes are logarithmic, lines parallel to this diagonal correspond to constant ratios of the two protein concentrations. Thus, the growth rate is observed to change only weakly when the protein concentrations are rescaled together by a common factor.

To uncover the origin of these observations, we switch from the protein-based description adopted in Fig 2, to a metabolite-based description (Secs. II C 2 and II E 2).

In the parametrization based on the concentrations of the metabolites, the growth time of the TR model is [eq. (48)]

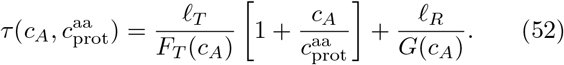

This objective function should be minimized with respect to its arguments. As the equality constraint (49c) is not present in the TR model, the minimization is subject only to the requirement that *c*_*A*_ and 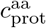 are non-negative. From the appearance of these variables in (52), it is clear that the growth time reaches its minimum in the limit 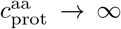. At the same time, from (45), the protein concentrations in the TR model satisfy

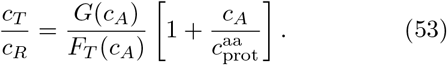

Thus, while the total protein concentration goes to infinity during the optimization of the growth rate, the ratio of the two protein concentrations remains finite. These observations explain the location of the maximum of the growth rate at the edge of the horizontal plane in Fig 2b.

Although the highest growth rate is reached for 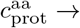 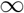, this limit is not physically meaningful. To offer an alternative way of interpreting it, we observe that the ratio 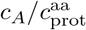 in (52) originates from the dilution of the metabolites on the left-hand side of (22). Thus, the limit 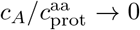 can alternatively be attained at finite protein concentrations but for vanishingly small metabolite concentration. Since any finite metabolite dilution, no matter how small, can only decrease the growth rate, the largest growth rate is achieved at negligible dilution of the metabolites.

### F. The role of metabolite dilution

The growth exhibited by our cell model (15) was a manifestation of the autocatalytic nature of the ribosomes (Sec. II A 3). The volume, in contrast, was needed only to convert the amounts of the metabolites to concentrations for use in the enzyme rate laws (Sec. II A 2). Because of this secondary role of the volume, we refrained from specifying its rate of change in (6), and subordinated its kinetics to the kinetics of the molecular amounts through the requirement of balanced growth [eq. (13)].

The central role of the ribosomes, however, was obscured by identifying the single kinetic degree of freedom that remained in balanced growth with the volume, since the other variables were retained only as constant multiples of it [i.e., the concentrations in (14)]. In this section, we select the amount of the ribosomes to be the reference degree of freedom. In the process, the special role played by the dilution of the metabolites is elucidated.

#### 1. The ribosome as a common reference

When the ribosome amount, *n*_*r*_, is used as a reference kinetic degree of freedom in balanced growth, the other kinetic variables are mapped to the following time-independent ratios:

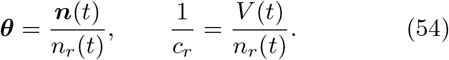

Notably, the ribosome concentration, *c*_*r*_, serves as a conversion factor between a normalization by *V* and a normalization by *n*_*r*_. Dividing our main equation (14) by this conversion factor, we get

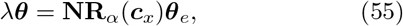

in which the variables are now normalized by *n*_*r*_, rather than the volume.

Similar to the partitioning of the vector ***n*** (Sec. II A), we split the vector of ratios, ***θ***, into its metabolic and enzyme subcomponents as follows:

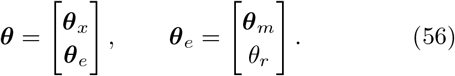

In the second equality, the enzyme ratios ***θ***_*e*_ are further split into the ratios of the metabolic enzymes, ***θ***_*m*_, and the ribosomes, *θ*_*r*_. Since *θ*_*r*_ = *n*_*r*_/*n*_*r*_ = 1, the vector ***θ***_*e*_ contains only *N*_*m*_ parameters.

Although the ratios ***θ*** reflect the central role of the ribosomes in our growth model, when it comes to the metabolites, it is more convenient to work with their concentrations since these appear in the rate laws. We therefore write the metabolic and protein parts of (55) as [*cf.* (15)]

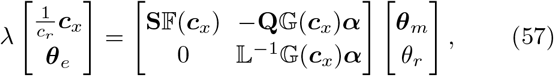

where the metabolites have been normalized by *V* and the enzymes by *n*_*r*_. The only place were the two different normalizations ever meet is the term *λ**c***_*x*_/*c*_*r*_, which is the product of the rate of metabolite dilution, *λ**c***_*x*_, and the volume expressed in ribosome units, *V/n*_*r*_ = 1/*c*_*r*_. Without this term, the two normalizations would be completely decoupled.

The difference between (55) and (14), or their expanded versions (57) and (15), consists of a simple rescaling. In that sense, (55) and (57) cannot be claimed to really bring forward the unique role of the ribosomes in the model. For this, one should look at the growth rate.

The growth time, which served as the objective function of the optimization problem (23), was given in (24). Here, we rewrite it in terms of the protein ratios, ***θ***_*e*_, as follows:

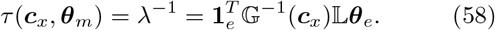

Since *τ* does not contain the concentration *c*_*r*_, which sets the scale of all other protein concentrations (***c***_*e*_ = *c*_*r*_***θ***_*e*_), we conclude that the growth rate in balanced growth does not depend directly on the absolute protein concentrations but only on their ratios with the ribosome concentration.

To make this last point explicit, we rewrite (58) as [*cf.*(47)]

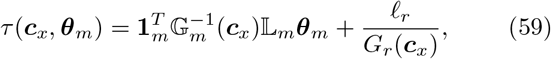

where 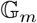and *G*_*r*_ are the parts of 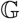 that correspond to the metabolic enzymes and the ribosomes, respectively. Clearly, this objective function is a function of *N*_*x*_ + *N*_*m*_ variables only (i.e., ***c***_*x*_ and ***θ***_*m*_), and not of *N*_*x*_ + *N*_*m*_ + 1 variables as suggested by (23b).

Of course, the ribosome concentration influences the growth rate indirectly through the metabolites, which are constrained by the upper part of (57). The corresponding constraint function is equal to ***h***_*x*_ [eq. (25)] divided by *c*_*r*_:

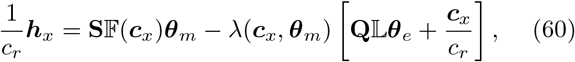

where *λ* is as given in (58). As noted before, all terms in (60) except the last one contain the enzyme ratios ***θ***_*e*_; the last term, which accounts for the dilution of the metabolites, depends on the concentration of the ribosomes.

At the end of Sec. II E 3, we observed that maximal growth rate is achieved when the entire mass flux is directed towards protein synthesis and the dilution of the metabolites is vanishingly small in comparison. Mathematically, this corresponds to the limit ***c***_*x*_/*c*_*r*_ → 0 in (60). In this limit, the concentration *c*_*r*_ disappears completely from the formalism. As a result, the normalization of the metabolites by the volume and the normalization of the proteins by the ribosomes decouple, and the absolute enzyme concentrations are unknowable.

#### 2. Neglecting metabolite dilution

In the absence of metabolite dilution, finding the root ***c***_*x*_ of the constraint function in (60) amounts to solving the equality

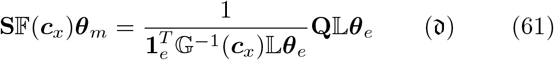

for the metabolite concentrations. (Equations that assume negligible metabolite dilution are indicated with 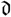.) Let ***c***_*x*_ be a solution of (61) for some protein concentrations. The same ***c***_*x*_ remains a solution after all protein concentrations are rescaled by a common factor, as such rescaling does not affect the ratios ***θ***_*e*_. Thus, when the metabolite dilution is neglected, the states of balanced growth depend only on the protein *composition* of the cell (i.e., the relative amounts of the proteins) and not on its protein content (i.e., the absolute protein concentrations). As the overall rescaling of the protein concentrations also leaves the growth rate of the model unaffected [eq. (58)], it appears that the protein concentrations can be increased indefinitely at fixed metabolite concentrations without affecting the growth rate.

To rationalize this flawed behaviour we note that the rate laws in (3) contain the concentrations of the metabolites, thus naturally coupling their amounts to the volume, but do not contain the *concentrations* of the enzymes, as only their amounts play a role. As far as the model is concerned, the enzymes might as well be point particles that do not occupy any volume. In reality, increasing protein concentrations will eventually force the volume to expand,^44^ providing direct coupling to volume that is not mediated by the dilution of the metabolites.

Although the formulated model lacks a built-in mechanism for keeping the enzyme concentrations reasonably low, its formulation actually assumed that this is the case. Indeed, the functional form of the enzyme rate laws applies only when the substrates are much more abundant than the enzymes. Additionally, increased macro-molecular crowding would progressively modify the rates of the modeled processes,^62,63^ rendering rate laws with constant catalytic parameters inappropriate. For the applicability of the mathematical analysis, therefore, we need to assume that the concentrations of the enzymes remain realistically small through some means that are not part of the model. (This limitation will be addressed in Sec. II H.) Subject to such a restriction, the solution ***c***_*x*_ of (61), which neglects metabolite dilution, is invariant to the simultaneous rescaling of the protein concentrations.

Naturally, the same conclusion applies to models with uniformly composed proteins (denoted by 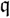 in Table 3), for which (61)simplifies to [*cf.* (44)]

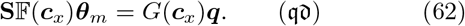

### G. Negligible metabolite dilution

The above observations clarify why the practical task of optimizing the growth rate was not addressed so far: if attempted such optimization would predict infinitely large protein concentrations. In this section we study the optimization of the growth rate assuming that the rate of metabolite dilution [*λ**c***_*x*_ in (22)] is negligible compared to the rates of production of the metabolites by the metabolic reactions 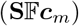 and their consumption in protein synthesis 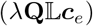.

#### 1. Optimization with the reduced gradient method

When the dilution of the metabolites is negligible, the constraint function in (23c) is replaced by [*cf.* (61)]

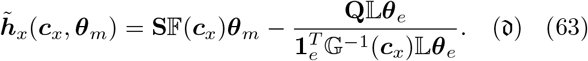

As both this constraint and the objective function [eq. (59)] are functions of the variables ***c***_*x*_ and ***θ***_*m*_, the optimization problem (23) becomes

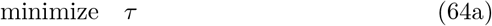

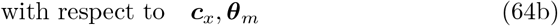

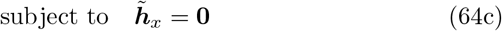

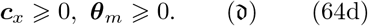

Since now *N*_*x*_ + *N*_*m*_ optimization variables are related through *N*_*x*_ equality constraints, only *N*_*m*_ variables are independent [compared to *N*_*m*_+1 = *N*_*e*_ variables in (23)].

With Δ***c***_*x*_ and Δ***θ***_*m*_ denoting small increments of the optimization variables (64b), the corresponding change in the objective function (64a) to first order in the increments is

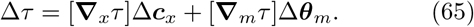

In this expression, the derivatives of the growth time are row vectors with components

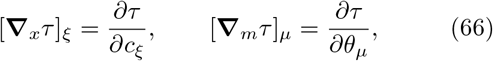

for *ξ* = 1, … , *N*_*x*_ and *μ* = 1, … , *N*_*m*_.

Naively, one could try to locate the minimum of the objective function by changing the optimization variables in the direction of its steepest descent. Such motion down the gradient of *τ*, however, is likely to violate the equality constraint (64c). A downhill direction that also satisfies the equality constraint, at least to first order in the increments, is given by the so-called reduced gradient.^64^

Assuming that the constraint was satisfied before the update of the variables, we require that it is also satisfied to first order after the update:^64^

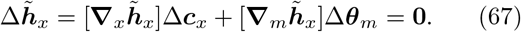

Here, the derivatives of the constraint function are matrices with components

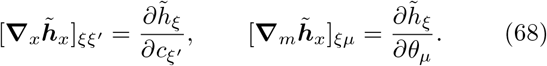

From (67), it is clear that ***c***_*x*_ and ***θ***_*m*_ cannot be changed independently while satisfying the constraint.

##### a. Proteins as independent variables

If the square matrix 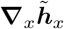 is invertible, (67) can be solved for Δ***c***_*x*_:

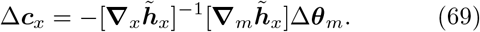

In Sec. S2B, we show that 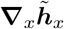 is invertible if the rate laws of the metabolic enzymes account for inhibition by their products. This is the reason why product inhibition was explicitly included in the rate laws of the metabolic reactions in Fig 1, even though the reactions were taken to be irreversible.

Substituting Δ***c***_*x*_ from (69) into (65), we obtain the reduced gradient of the objective function with respect to the independent protein variables:^64^

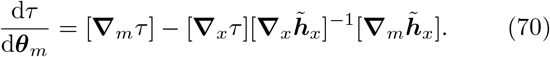

This gradient is a row vector with *N*_*m*_ components.

The independent protein variables are now updated in the direction opposite to the reduced gradient:^64^

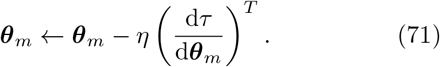

The step size *η* should be small enough for the value of the objective function at the new point to be smaller than its current value.

Although the corresponding increments of the dependent metabolite variables can be calculated from the increments of the independent variables using (69), this proportionality is only accurate to first order. A better strategy is to update the dependent variables using either the Newton-Raphson algorithm (26) or the multiplicative update (28), but with neglected metabolite dilution.

##### b. Metabolites as independent variables

For completeness, we also mention the possibility of using the metabolite concentrations, ***c***_*x*_, as the independent variables of the optimization problem (64). As discussed in Sec. II C 2, this option was available only for models with **S** of full column rank (denoted by 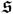 in Table 3). When the number of metabolites was larger than the number of metabolic enzymes (*N*_*x*_ > *N*_*m*_), the *N*_*x*_ equality constraints (23c) led to the *N*_*m*_ equalities (32) and the *N*_*x*_ − *N*_*m*_ conservation equalities (34).

When the dilution of the metabolites is neglected, as done presently, the metabolite concentrations, ***c***_*x*_, drop out completely from the conservation relations (34), thus compromising these constraints. To avoid such complications, we will illustrate the possibility of treating the metabolites as the independent variables of the optimization (64) only for the case with equal numbers of metabolites and metabolic enzymes (*N*_*x*_ = *N*_*m*_). As **S** was assumed to be of full column rank, this condition makes it invertible, and does away with the conservation relations (34).

To express the increments Δ***θ***_*m*_ in (67) in terms of the increments Δ***c***_*x*_, the *N*_*x*_ × *N*_*m*_-matrix 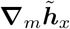 in (67) should be “inverted”. When *N*_*x*_ = *N*_*m*_ and **S** is invertible, 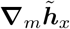 is also invertible if none of the diagonal elements of 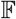 is equal to zero (as discussed in the Sec. S2B). The reduced gradient of the objective function with respect to the independent metabolite variables is now

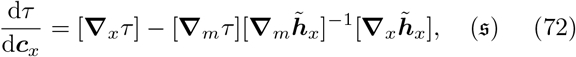

and the concentrations of the metabolites are updated as

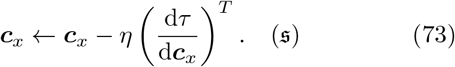

At every time step, the dependent variables ***θ***_*m*_ can be calculated using the multiplicative update rule (35), but with neglected metabolite dilution.

#### 2. Numerical examples

##### a. The TR model

We repeated the analysis of the TR model but with the last term of *h*_*A*_ in (50) set to zero (i.e., neglecting metabolite dilution). As expected, the contours of con-stant *c*_*A*_ (Fig 3a) as well as those of constant *λ* (Fig 3b) are now perfectly aligned with the *c*_*T*_ = *c*_*R*_ diagonal of the protein plane, thus illustrating the invariance of the balanced-growth states to the overall rescaling of the protein concentrations. Clearly, when the dilution of the metabolites is neglected, the ratio *θ*_*T*_ = *c*_*T*_/*c*_*R*_ is sufficient to parametrize the states of balanced growth.

**Fig 3.**
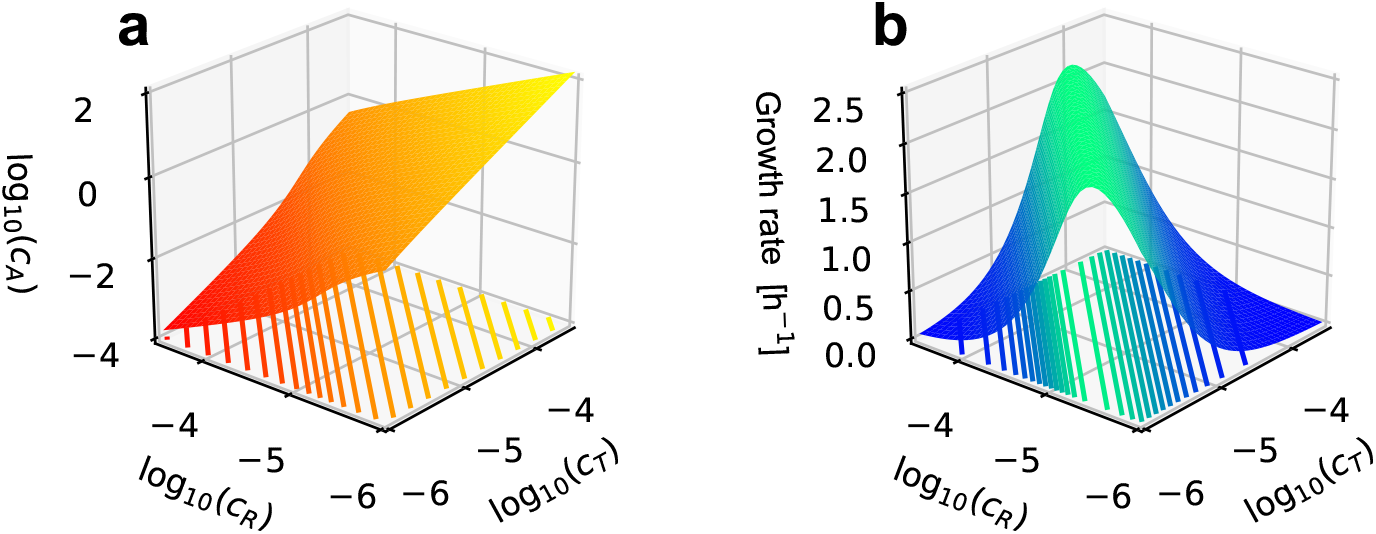
TR model as in Fig 2, but with neglected metabolite dilution.

In Fig 4a we show the metabolite concentration (left vertical axis) and the growth rate (right vertical axis) of the TR model against this parameter. The two curves are the cross-sections of the respective surfaces in Fig 3. A comparison with the Michaelis constants of the two proteins reveals that the inflection points of the *c*_*A*_ curve (red solid line) correspond to the values of *K*_*T*_ and *K*_*R*_ (black dashed lines). In this particular case, the growth rate is seen to be maximal when *c*_*A*_ ≈ *K*_*T*_, at which point the ribosome works at close to full saturation.

**Fig 4.**
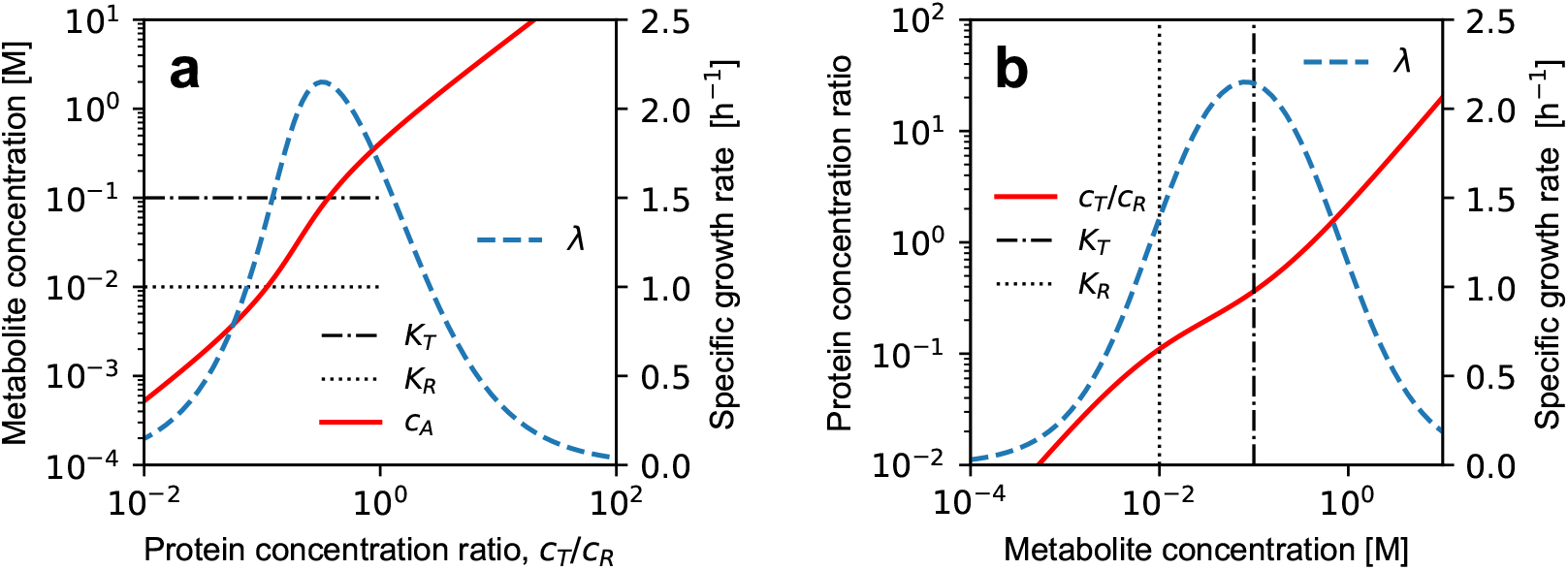
Analysis of the TR model with neglected metabolite dilution. The states of balanced growth are parametrized by either the ratio of the two protein concentrations (**a**) or the metabolite concentration (**b**).

For the TR model with neglected metabolite dilution, the equality constraint (62) is

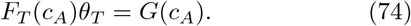

Because the functions *F* and *G* are nonlinear in their argument, this equality had to be solved for ***c***_*A*_ numerically when generating Fig 4a. In contrast, (74) it is trivial to solve for *θ*_*T*_, assuming a given *c*_*A*_:

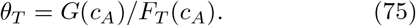

Using this expression, we calculated the ratio *c*_*T*_/*c*_*R*_ for *c*_*A*_ between 0.1 *μ*M and 10 M. The result, together with the corresponding growth rate, is shown in Fig 4b. Although for the red curve this is just Fig 4a with swapped horizontal and vertical axes, no iterative updates were necessary for its generation.

To numerically locate the optimal growth rate of the TR model, we applied the reduced gradient algorithm treating either the protein ratio, *θ*_*T*_, or the metabolite concentration, *c*_*A*_, as the independent variables. At convergence, we had *λ* = 2.15 h^−1^, *θ*_*T*_ = 0.324 and *c*_*A*_ = 81.7 mM in both cases. These values agree visually with the locations of the maximal growth rate in Figs 4a and 4b.

##### b. The TVR model

In the case of the TVR model with neglected metabolite dilution, the ratios *θ*_*T*_ = *c*_*T*_/*c*_*R*_ and *θ*_*V*_ = *c*_*V*_/*c*_*R*_ parametrize all balanced-growth states. Because there is only one metabolite, there is only one equality constraint. Figure 5a shows the metabolite concentration, located with the Newton-Raphson algorithm, against the ratios *θ*_*T*_ and *θ*_*V*_. The corresponding growth rate is plotted in Fig 5b. Identical results were obtained using the multiplicative update (28) with *κ* = 1:

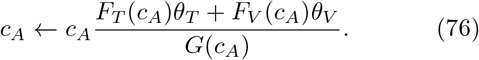

**Fig 5.**
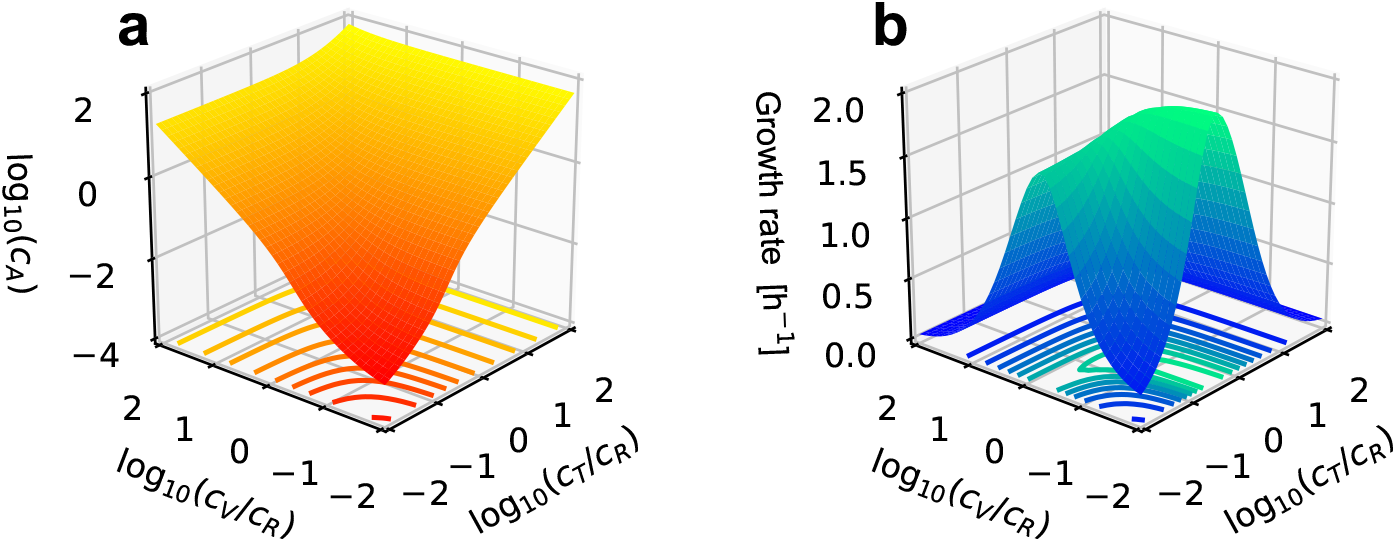
Balanced growth analysis of the TVR model with neglected metabolite dilution. Parametrization by the ratios of the concentrations of the metabolic proteins to the concentration of the ribosome.

When the concentration of either *T* or *V* relative to that of the ribosome is vanishingly small, the TVR model reduces to the TR model. Indeed, both the *θ*_*T*_ = 10^−2^ and *θ*_*V*_ = 10^−2^ cross-sections of the surface in Fig 5b resemble the profile of the growth rate in Fig 4a. Due to our choice of enzyme parameters (Table 2), the maximal growth rate for *θ*_*T*_ = 10^−2^ is smaller than that for *θ*_*V*_ = 10^−2^. Hence, the latter is the global maximum, at least for the range of *θ*_*T*_ and *θ*_*V*_ values shown in the figure.

For the TVR model, the equality constraint (62) reads

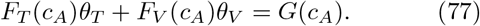

While (77) could be solved for *c*_*A*_ when *θ*_*T*_ and *θ*_*V*_ were given, the opposite, i.e., calculating these two protein ratios for given *c*_*A*_, is not uniquely possible. From that perspective, the TVR model is different than the TR model. Indeed the metabolic stoichiometry matrix of the former is not of full column rank and does not have a left inverse (it is of class 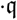 in Table 3). Thus, the balanced-growth states of the TVR model cannot be fully parametrized in terms of the metabolites rather than the ratios of the protein concentrations.

Because the growth rate attains its maxima when either *θ*_*V*_ → 0 (global maximum) or *θ*_*T*_ → 0 (local maximum), during the optimization of the growth rate it is advantageous to work with the square roots of *θ*_*T*_ and *θ*_*V*_ in order to prevent the vanishing ratio from becoming negative in the update (71). Applying the reduced gradient method we obtained the optimal growth rate *λ* = 2.15 h^−1^ at *θ*_*T*_ = 0.324, *θ*_*V*_ = 0.00 and *c*_*A*_ = 81.7 mM. This corresponds to the global maximum in Fig 5b. Only when the search was initiated very near the second (local) maximum did the algorithm converge to *λ* = 1.68 h^−1^ at *θ*_*T*_ = 0.00, *θ*_*V*_ = 0.532 and *c*_*A*_ = 46.3 mM.

##### c. The TUR model

As the matrix **S** of the TUR model is invertible (it is of class 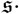 in Table 3), its balanced-growth states allow full parametrization in terms of the metabolites. From these, the protein ratios can be determined iteratively using the multiplicative update (36), after dropping the contribution of metabolite dilution. For *κ* = 1, and *θ*_*T*_ = *c*_*T*_/*c*_*R*_ and *θ*_*U*_ = *c*_*U*_/*c*_*R*_, this update reads

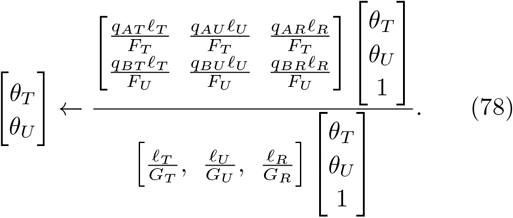

At convergence, the denominator of (78) is equal to the growth time, *τ*.

Both surfaces *θ*_*T*_ and *θ*_*U*_ (Fig 6a) converged to five (ten) decimal places in 10 (20) iteration updates across the entire range of examined metabolite concentrations.(These were calculated for the *k*_cat_ and *K*_*M*_ values in Table 2.) As expected, *θ*_*T*_ depends strongly on the concentration of *A* but only weakly on the concentration of *B*. In contrast, *θ*_*U*_ increases with increasing *c*_*B*_ but is almost insensitive to *c*_*A*_.

**Fig 6.**
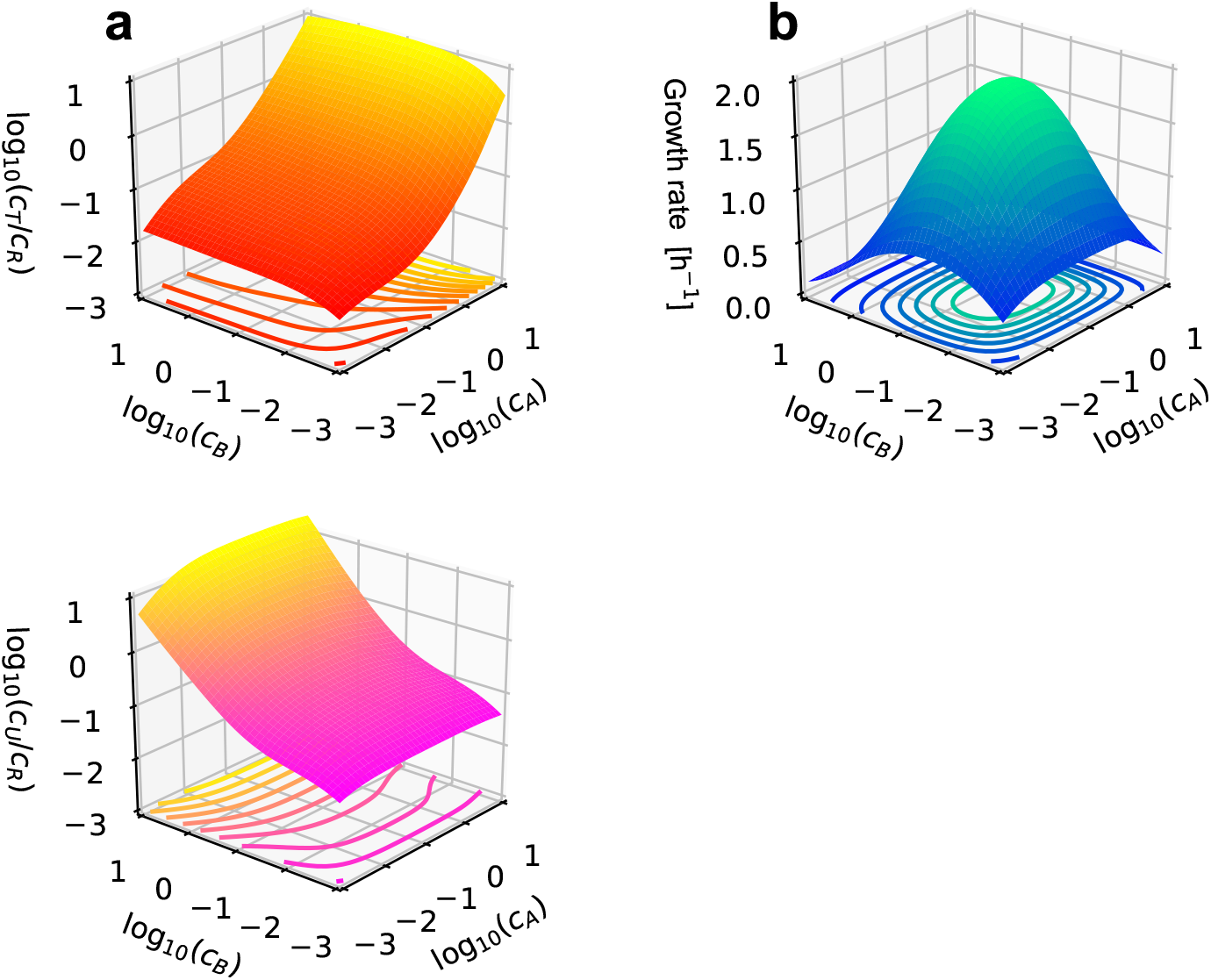
Balanced growth analysis of the TUR model with neglected metabolite dilution. The ratios of the metabolic proteins to the ribosome (**a**), and the growth rate (**b**), are parametrized by the concentrations of the metabolites.

The growth rate in Fig 6b reaches its global maximum in the middle of the examined range of metabolite concentrations. Unlike the situation in the TVR model (Fig 5b), here none of the proteins vanishes at the maximal growth rate, as both are required for cell growth.

In the optimization of this model, either the protein ratios or the metabolite concentrations can be treated as independent variables. To be consistent with Fig 6, we made the latter choice when applying the reduced gradient method. Consequently, the independent concentrations *c*_*A*_ and *c*_*B*_ were updated according to (73) while the dependent protein ratios *θ*_*T*_ and *θ*_*V*_ were determined using the multiplicative update (78). The algorithm converged to *λ* = 2.03 h^−1^ at *c*_*A*_ = 83.1 mM and *c*_*B*_ = 37.7 mM, with corresponding protein ratios *θ*_*T*_ = 0.099 and *θ*_*U*_ = 0.270. This point agrees with the maximal growth rate in Fig 6b.

### H. Constraining the cell volume

Metabolite dilution was neglected in the previous section for purely mathematical reasons: if accounted for, it forced the enzyme concentrations to increase indefinitely during the optimization of the growth rate. By neglecting it, however, we effectively took the stance that the enzyme concentrations were fundamentally unknowable and only their ratios were meaningful. Ultimately, this position is indefensible.

The origin of the problem is addressed in the current section by assigning molar volumes to the enzymes of the model. Consequently, protein concentrations become limited, and the optimal growth rate is reached at finite protein concentrations, even after accounting for the dilution of the metabolites.

#### 1. The volume in balanced growth

By leaving the rate of change of the volume unspecified in (6), we gave up on the possibility to model the full kinetics of the dynamical system comprising eqs. (5) and (6). The latter equation, together with the condition of balanced growth [eq. (13)], implies

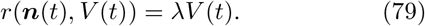

In principle, this equality can be solved for the volume, thus expressing *V* in balanced growth as a function of the molecular amounts and the growth rate:

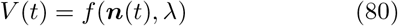

Formally, this functional dependence can be used to introduce the partial molar volumes of the substances at a given growth rate:

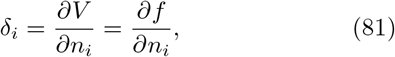

where *δ*_*i*_ is the change in volume per mole of substance *i* added to a large volume of the mixture.^65^

On the other hand, differentiating (80) with respect to time, and using the conditions of balanced growth (13), we get

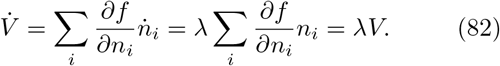

Combining the last equality in (82) with the definition of the partial volumes in (81), we deduce that the volume in balanced growth is a weighted sum of the amounts of the cell constituents:^65^

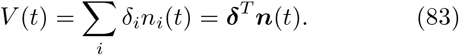

In mixtures of non-interacting components, the partial molar volumes, ***δ***, are equal to the molar volumes of the components. When the components interact, however, the partial molar volumes not only deviate from the molar volumes but may even be negative.^65^ Since the contents of the cell likely behave as an interacting mixture, the *N*_*x*_ + *N*_*e*_ partial molar volumes in (83) are not expected to be known precisely in any specific situation. Nonetheless, a vast majority of these coefficients will certainly be strictly positive.

The importance of (83) for our purposes is that it assigns volumes to the proteins of the model. As a result, their amounts cannot be increased indefinitely without increasing the cell volume. By providing a direct coupling of the enzymes to the volume, as opposed to the indirect coupling through the dilution of the metabolites, the model now has an in-built mechanism for restricting the concentrations of the proteins. What remains to be done is to turn (83) into a practical tool for the mathematical analysis, even if the partial molar volumes are not known.

#### 2. Constant density approximations

Measurements of the cytoplasmic protein and RNA content of the bacterium *E. coli* revealed that the densities of these two major mass components varied across exponential and stationary growth phases with their total density remaining approximately constant.^66^ A more recent experimental study reported the proportionality of volume and cell dry mass for exponentially growing *E. coli* under various growth limitations,^67^ suggesting that the dry mass density is approximately constant. Whereas the older reported values of the total cytoplasmic protein and RNA density were spread between 300 g l^−1^ and 400 g l^−1^,^66^ recent optical measurements indicate that the dry mass density of *E. coli* varies more narrowly between 270 g l^−1^ and 320 g l^−1^ depending on the growth medium.^68^

By definition of the dry mass density, *ρ*_dry_ (g l^−1^), the relationship between the cell volume and the amounts of the cell constituents, excluding water, is

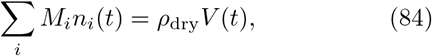

where *M*_*i*_ (g mol^−1^) are the molar masses of the chemical species. Dividing both sides of (84) by the volume, the dry mass density is readily expressed in terms of the concentrations of the species:

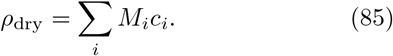

To calculate *ρ*_dry_ of the TR model, we assigned to its metabolite *A* the molar mass *M*_*A*_ = 100 g mol^−1^, which corresponds to an average amino acid.^58^ Then, from (85), the dry mass density of the model is *ρ*_dry_ = *M*_*A*_(*c*_*A*_ + *ℓ*_*T*_*c*_*T*_ + *ℓ*_*R*_*c*_*R*_). In Fig 7, the lines of constant *ρ*_dry_ are plotted on top of the contours of the growth rate from Fig 2b. Although the growth rate in the figure is calculated for all pairs of protein concentrations, the above discussion suggests that only the narrow vicinity of the contour *ρ*_dry_ = 300 g l^−1^ should be viewed as biophysically relevant. Clearly, if the optimization of the growth rate is restricted to remain on this contour, the protein concentrations at the optimal growth rate will be finite.

**Fig 7.**
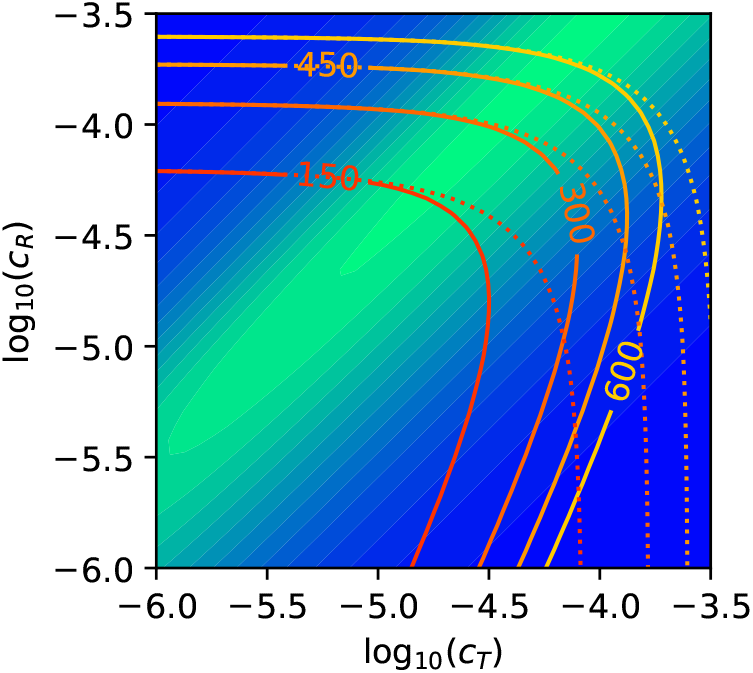
Contours of constant growth rate (filled color, identical to the contours in Fig 2b), and the densities *ρ*_dry_ (solid, g l^−1^) and *ρ*_macro_ (dotted) of the TR model.

In addition to the catalytic proteins, the ribosomes, and the metabolites, which are represented in our illustrative models, the dry mass of the cell receives contribution from structural proteins, genomic DNA, lipids, lipopolysaccharides, peptidoglycan, and other types of molecules, whose functional roles are not part of these models. As such structural and informational molecules could constitute a substantial fraction of the cell dry mass, the experimental density should, in principle, be rescaled when used in these models. Given the coarse-grained nature of the models, however, it is not clear what exactly the correct rescaling should be. Since the examples in this section are intended to serve as qualitative illustrations, we have not performed any rescaling.

One way of enforcing a constant dry mass density through eq. (83) is to select *δ*_*i*_ = *M*_*i*_/*ρ*_dry_ for all *i*. This corresponds to a mixture of non-interacting components, whose molar volumes are proportional to their molar masses.

Alternatively, the cytoplasm could be envisioned as a highly interacting mixture whose volume is dominated by the molar volumes of the hydrated macromolecules, while the comparatively smaller metabolites occupy the space between them without requiring too much additional hydration, thus having partial molar volumes that are close to zero [i.e., large ***δ***_*e*_ and ***δ***_*x*_ ≈ 0 in (83)]. In this case, the assumption of constant dry mass density can be replaced by an assumption of (approximately) constant macromolecular density. Since all macromolecules in our model are proteins, this assumption can be stated as

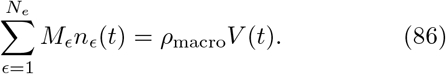

Dividing both sides of (86) by the volume, the macro-molecular density in our model is expressed in terms of the concentrations of the enzymes:

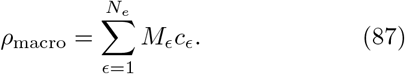

As an example, the lines of constant *ρ*_macro_ of the TR model are shown in Fig 7. Naturally, they are almost identical to the contours of *ρ*_dry_ in the upper left corner of the figure, where *c*_*A*_ is small (*cf.* Fig 2a), but deviate substantially in the lower right corner where *c*_*A*_ is large.

One way of expressing the assumption of constant macromolecular density through eq. (83), is to select *δ*_*ϵ*_ = *M*_*ϵ*_/*ρ*_macro_ for all enzymes and ***δ***_*x*_ = 0 for the metabolites. Taken literally, this choice implies a non-interacting mixture composed of macromolecules, whose molar volumes are proportional to their molar masses, and metabolites with negligibly small volumes.

Whether any of the densities *ρ*_dry_ and *ρ*_macro_ is indeed approximately constant across different balanced-growth conditions is an experimental question. However, considering that the metabolites constitute about 5-10% of the dry mass of *E. coli* cells,^58,60^ the distinction between the dry mass density and the macromolecular density may be inconsequential for the purpose of constraining the enzyme concentrations during the optimization of the growth rate.

#### 3. Optimization with constrained density

Neglecting the metabolite dilution term *λ**c***_*x*_/*c*_*r*_ from the constraint function (60) allowed us to carry out the optimization (64) in the space of the metabolite concentrations, ***c***_*x*_, and the ratios of the metabolic enzymes, ***θ***_*m*_. To bring metabolite dilution back into the optimization procedure of Sec. II G 1, it is sufficient to express the ribosome concentration, *c*_*r*_, in terms of the optimization variables (64b).

Dividing both sides of (83) by the amount of the ribosomes, *n*_*r*_, we get

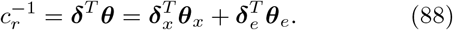

Here, both the metabolites and the enzymes appear as the ratios ***θ*** [*cf.* (56)]. Since it is more convenient to work with the concentrations of the metabolites, rather than their ratios, we use 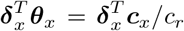 in (88) to deduce that

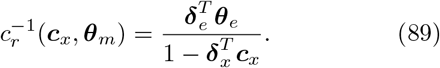

This is a closed-form expression of the reciprocal of the ribosome concentration in terms of the optimization variables (64b).

##### a. Constant macromolecular density

The assumption of constant macromolecular density corresponds to using ***δ***_*x*_ = 0 for the metabolites and *δ*_*ϵ*_ = *M*_*ϵ*_/*ρ*_macro_ for the enzymes in (89). With this choice, the concentration of the ribosomes is a function of the enzyme ratios only:

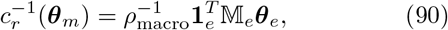

where the matrix 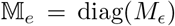, *ϵ* = 1, … , *N*_*e*_, contains the molar masses of the enzymes. The analysis of Sec. II G 1 is thus directly applicable after subtracting the following extra term from the constraint function 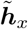 in (63):

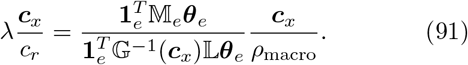

(The derivatives of this new constraint function with respect to the metabolite concentrations, ***c***_*x*_, and the ratios of the metabolic proteins, ***θ***_*m*_, are given in Sec. S3 of the SI.)

For models with uniformly composed proteins, 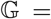 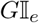 and 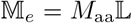, where *M*_aa_ (g mol^−1^) is the molar mass of an average amino acid. Then, (91) simplifies to

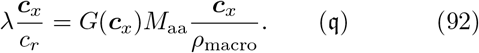

In the case of the TR and TVR models (class 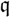 in Table 3), the equality constraints in (74) and (77) now become

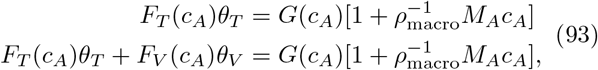

respectively. Consequently, the functions *G* in (75) and (76) should be multiplied by the factor 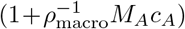.

In the case of the TUR model, the following extra contribution should be added to the numerator of the multiplicative update (78):

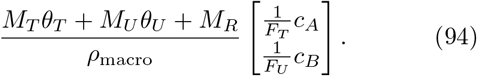

Its denominator remains unmodified.

The optimization of these three illustrative models was carried out with the reduced gradient method after introducing the outlined changes. We used *ρ*_macro_ = 300 g l^−1^, and *M*_*A*_ = *M*_*B*_ = 100 g mol^−1^ for the molar masses of the metabolites. The results are compiled in rows ‘m’ (for macro) of Table 4. The previous results with neglected metabolite dilution were recovered using 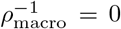 in the above equations (rows ‘n’ of Table 4). As this latter case leads to effectively infinite protein concentrations, the concentration of the ribosome was left unspecified in the last column of the table.

**Table 4.** Optima of the three illustrative models, located with neglected metabolite dilution (n), assuming constant macro-molecular density *ρ*_macro_ = 300 g l^−1^ (m), or constant dry mass density *ρ*_dry_ = 300 g l^−1^(d).

| | | $\lambda/\text{h}^{-1}$ | $\theta_T$ | $\theta_{V,U}$ | $c_A/\text{mM}$ | $c_B/\text{mM}$ | $c_R/\mu\text{M}$ |
| --- | --- | --- | --- | --- | --- | --- | --- |
| TR | n | 2.1506 | 0.3237 | - | 81.65 | - | - |
|  | m | 2.1396 | 0.3246 | - | 78.37 | - | 100.53 |
|  | d | 2.1393 | 0.3245 | - | 78.23 | - | 97.91 |
| TVR | n | 2.1506 | 0.3237 | 0.0000 | 81.65 | - | - |
|  | m | 2.1396 | 0.3246 | 0.0000 | 78.37 | - | 100.53 |
|  | d | 2.1393 | 0.3245 | 0.0000 | 78.23 | - | 97.91 |
| TUR | n | 2.0272 | 0.0992 | 0.2703 | 83.14 | 37.70 | - |
|  | m | 2.0117 | 0.1011 | 0.2699 | 73.76 | 36.47 | 95.27 |
|  | d | 2.0111 | 0.1010 | 0.2698 | 73.25 | 36.40 | 91.80 |

##### b. Constant dry mass density

In the case of constant dry mass density, we have 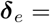 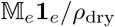 for the enzymes as well as 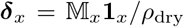 for the metabolites, where the molar masses of the latter have been collected in the diagonal matrix 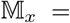 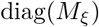, *ξ* = 1, … , *N*_*x*_. Substituting these partial molar volumes into the expression of the ribosome concentration (89), we get

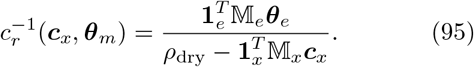

The derivatives of the resulting constraint function with respect to ***c***_*x*_ and ***θ***_*m*_ are given in Sec. S3B.

The optimization results for the three illustrative models are given in rows ‘d’ (for dry mass) of Table 4. A comparison of the optimal growth rates in the ‘n’, ‘m’ and ‘d’ rows of each model reveals that, for the selected model parameters, the additional constraint on the density reduces the growth rate by less than 1%. In the case of the TR model, this result is in line with Fig 7, where the contours of constant growth rate show that the maximal *λ* has plateaued out already at densities of about 150 g l^−1^. Thus, increasing the density from 300 g l^−1^ to 600 g l^−1^ or all the way to infinity (row ‘n’) is not expected to significantly change the maximal growth rate.

Similar to the growth rate, the protein fractions at the optimum are also observed to depend only weakly on the density constraint. The largest effect, which is still moderate, is on the concentrations of the metabolites.

From the perspective of the current section, the observed weak effect of the density constraint on the optimal growth rate and the corresponding protein ratios (though not necessarily on the metabolite concentrations) is extremely encouraging. After all, the whole idea of fixing either the dry mass density or the macromolecular density (and not only its precise numerical value) is a convenient shortcut that, ideally, should be replaced by a more realistic model of cell size regulation.

##### c. Neglected metabolite dilution

The constraint on the density was introduced above in order to account for the dilution of the metabolite concentrations in the optimization of the growth rate. Nevertheless, the physical picture behind the relationship between cell volume and molecular amounts [eq. (83)], as well as the assumptions that led to eqs. (90) or (95) for the ribosome concentration, should be largely independent of the magnitude of the dilution term. The same equations are therefore expected to hold even when the dilution of the metabolites is negligibly small. Then, it becomes possible to maximize the growth rate in the absence of metabolite dilution, as was done in Sec. II G, and subsequently calculate *c*_*r*_, using either (90) or (95), to recover finite protein concentrations at the optimum. When applied to the TUR model, for example, this approach yields either *c*_*r*_ = 95.35 *μ*M (with *ρ*_macro_ = 300 g l^−1^) or *c*_*r*_ = 91.51 *μ*M (with *ρ*_dry_ = 300 g l^−1^). These values, calculated after an optimization with neglected metabolite dilution, are intended to replace the missing ribosome concentration in row ‘n’ of Table 4 for the TUR model.

## III. Discussion

### A. Summary

One of the main aims of the current paper was to bridge the gap between the analyses of refs. 31 and 32, both of which addressed balanced cell growth in a general way applicable to large, genome-scale models. Here, we clarified that the principal difference between the formalisms of these two studies is the choice of independent variables used to parametrize the states of balanced growth—metabolite concentrations in the case of the former and ribosome allocation fractions in the latter (Sec. II E). In the process, the formalism was developed for yet another choice of these variables: the concentrations of the proteins.

The other central goal of the paper was to bridge the gap between the analyses of refs. 31 and 32, on the one hand, and the actual numerical optimization of the growth rate in balanced growth, on the other. In Sec. II C, we showed how to numerically calculate the concentrations of the metabolites from known protein concentrations within the framework of a kinetic, autosynthetic cell model of balanced growth. To this end, multiplicative updates were introduced as an alternative to Newton-Raphson’s additive updates, thus avoiding the need for the derivatives of the rate laws and the inversion of the Jacobian matrix. Multiplicative updates appear to be perfectly suited to the current problem since reaction fluxes can always be separated into positive (production or inflow) and negative (consumption or outflow) parts.^69^

Given the more extensive experimental coverage of the cell proteome (e.g., 2359 protein concentrations of *E. coli* were measured under 22 experimental conditions^70^) in comparison to the cell metabolome (103 metabolite concentrations under 3 conditions^60^), the proposed computational framework could, in principle, be useful for predicting the metabolite concentrations that correspond to a given proteome. In practice, the bottleneck in this task will be poor knowledge of the enzyme rate laws and their kinetic and thermodynamic parameters.^15^ However, efforts for expanding the existing *k*_cat_ and *K*_*M*_ values^16,17^ by machine-learning approaches are underway.^71^ Provided these parameters are available, the reduced gradient method of Sec. II G 1 should allow for the practical numerical optimization of the growth rate of large, autosynthetic cell models.

In addition to the aforementioned primary objectives, some issues pertaining to the states of balanced growth, in general, and to the problem of optimizing the balanced growth rate, in particular, were also addressed.

In Sec. II F, the growth-induced dilution of the metabolite concentrations was identified to be the only point in the treatment where the amounts of the enzymes coupled to the volume of the cell. This is because, on the one hand, the amounts of all chemical species are ultimately proportional to the amount of the ribosomes and, on the other, only the metabolites are “perceived” (through the enzyme rate laws) as concentrations rather than as absolute amounts. By simultaneously “sensing” both the volume of the cell and the ribosome amount, the metabolite dilution provides a weak, indirect coupling between the two. Although the magnitude of this term influences the growth rate, maximal growth rate is achieved at vanishingly small metabolite dilution, which corresponds to infinite protein concentrations.

In the light of this observation, two alternative ways of dealing with the problem of infinite protein concentrations were subsequently explored. In Sec. II G, we neglected the dilution of the metabolites and observed that the resulting model behaves well (i.e., problems with matrix inversion are avoided) if the inhibition of the enzymes by their products is accounted for in the rate laws. This mathematical requirement highlights the mechanistic role that product inhibition plays in restricting the metabolite concentrations in the cell. Indeed, the concentrations of some metabolites could become unrealistically high in models with missing product inhibition. In these cases, including the metabolites in the density constraint serves as a natural constraining mechanism.

In Sec. II H, the dilution of the metabolites was reinstated after directly linking the cell volume to the amounts of the proteins by assuming constant macro-molecular density or dry-mass density. This direct coupling trumped the indirect coupling mediated by the dilution of the metabolites, and forced the concentrations of the proteins to remain finite during the optimization of the growth rate.

The discussion in Sec. II H 1, as well as physical intuition, are unequivocal about the direct effect that cell constituents exert on cell volume. In this context, the assumption of constant dry mass density is frequently invoked as an integral part of models of balanced cell growth.^41^ In particular, the dry mass density was constrained in both refs. 31 and 32.

While the mathematical role of the assumption of constant density for the optimization of the growth rate is unquestionable, the numerical examples in Sec. II H 3 suggest that envisioning the density as a limited physical resource for which the (macro)molecules have to compete may be misleading in some cases. In the considered examples, the optimal states of the three models with constrained dry mass or macromolecular density were practically identical to the states at infinite density (i.e., with neglected metabolite dilution). The density constraint thus appears to only perturb the optimal state, rather than dictate it.

Of course, the cell density may be a stiff and dominant constraint under some growth conditions and a soft constraint under others. The observation that protein synthesis continued even after cell growth was inhibited in fission yeast is an example for the latter.^44^

### B. Relation to other formulations

Both refs. 31 and 32 analyzed a cell model that grows by synthesizing its own enzymes. This problem has been studied before both in the context of specific small-scale models^26–29,72,73^ and more generally.^20,21,33,74,75^ However, a closer examination of the constrained optimization problems formulated in these and other similar studies often reveals subtle differences in either the choice of the objective function or the mathematical expressions of the constraints. Sometimes these variations make it hard to decide which conclusions depend on the particular choices made and which apply more generally to other optimization scenarious.

For example, both refs. 31 and 32 include the metabolites in their density constraints. In contrast, only the total density of the enzymes (i.e., the macromolecules of the model) is constrained in refs. 28, 29, and 33. Another common difference consists in the treatment of the metabolite concentrations: their dilution by growth is either included^31,32,73^ or omitted.^21,29,75^ Yet another point of variation is the choice of the objective function. While we and others^21,26–29,31–33,72,73^ have selected to maximize the growth rate of the model at a fixed density (either *ρ*_dry_ or *ρ*_macro_), some studies minimize the total enzyme density at a fixed output flux (e.g., the flux through the “ribosome” in the case of growth).^19,20,75^

In Fig 8, the optimization studied in the current paper is depicted as being at the top of a hierarchy of possible optimization problems that are reached after one or two simplifying assumptions. The scheme we examined is in the rectangle at Level 0. By neglecting the dilution of the metabolite concentrations, we descend one level down (Level 1). This assumption is commonly invoked.^20,21,29,75^ Further down (Level 2) are optimization problems that assume all enzymes to operate with metabolite-independent molar rates. Resource balance analysis (RBA) models^33,34,74^ fall into this last category.

**Fig 8.**
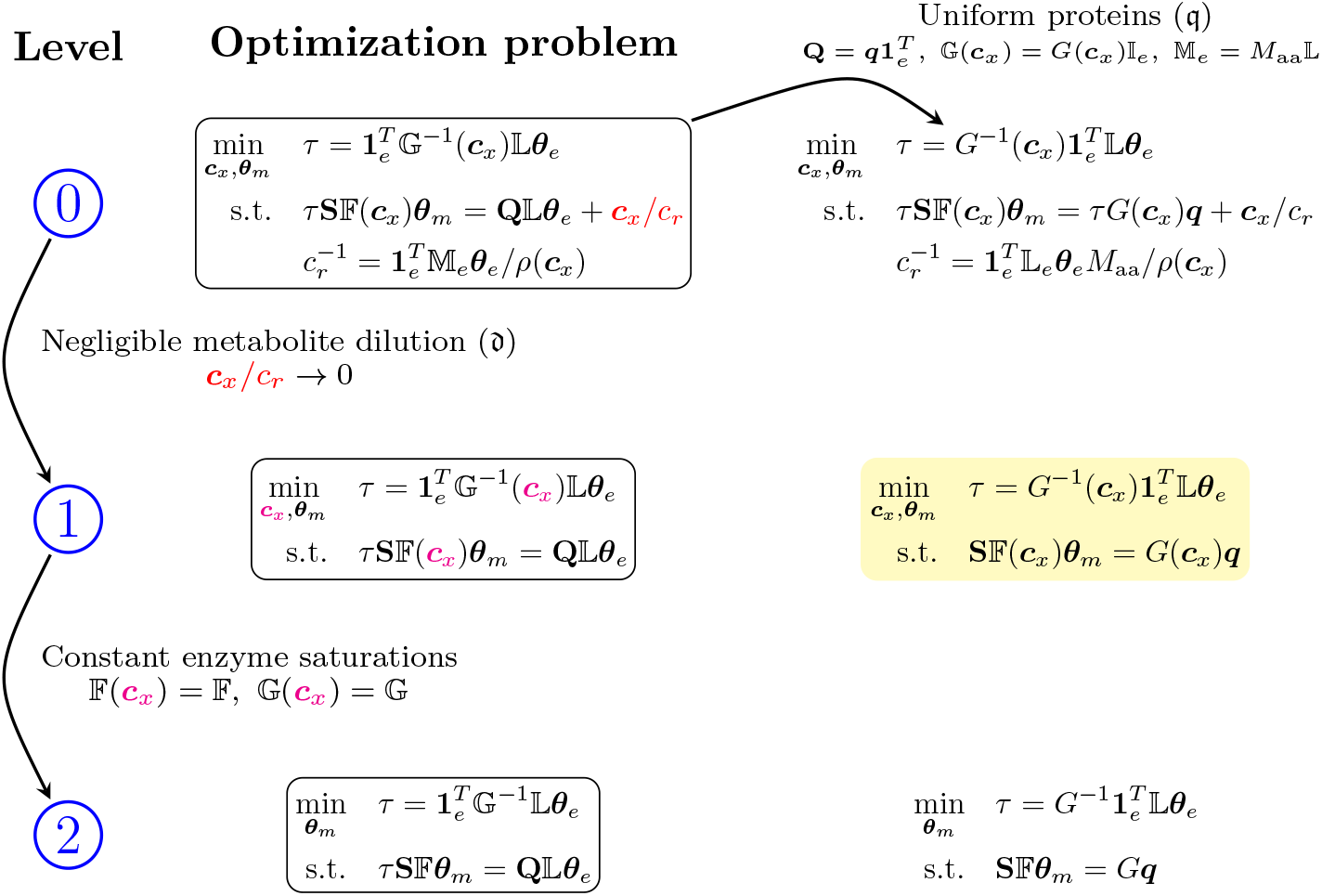
A hierarchy of growth-rate optimization problems resulting from (i) neglecting the dilution of the metabolite concentrations, denoted by 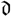, and (ii) additionally assuming that the saturations of all enzymes remain constant. A parallel hierarchy exists for models whose proteins have identical amino acid composition, which were denoted by 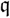. Most of the general analyses in the literature are related to the optimization highlighted in yellow.

Given that the metabolites are missing entirely from Level 2, the question arises as to what exactly the optimization of the growth rate achieves here. An answer is given in ref. 34, which offers a concise but very readable account of RBA. At this level, the optimization decides which reactions should be active and which should be inactive. The ability to switch parts of the metabolic network on and off allows for, among others, modeling the transition to fermentation in overflow metabolism and the hierarchical utilization of various carbon sources in catabolite repression.^34^

Building on this background, the optimization at the higher level (Level 1) additionally determines the metabolite concentrations that maximize the fluxes of the active reactions by accounting for the dependence of the enzyme saturations on the metabolites. This information may feed back on the level below to change the ranking of two alternative active subnetworks, as has been illustrated on a simple example in ref. 21. At the highest level (Level 0), the concentrations in the active subnetwork are further modified by taking into account the contribution of metabolite dilution.

For the optimization at Level 0, a constraint on the density is needed in order to express the concentration of the ribosomes in terms of the optimization variables ***c***_*x*_ and ***θ***_*m*_ (Sec. II H). In Fig 8, the density constraint is present in the function *ρ*(***c***_*x*_), which is equal to either *ρ*_macro_ [eq. (90)] or 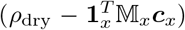 [eq. (95)], though other choices for the relative weights of the different molecular species are also conceivable.^32^ At the two lower levels of the hierarchy (Levels 1 and 2), the assumption of constant density is not necessary to carry out the optimization because the ribosome concentration drops out together with the neglected metabolite dilution. If desired, such a constraint can be employed after the optimization to calculate the concentration of the ribosomes and, subsequently, the concentrations of the metabolic enzymes (Sec. II H 3 c).

The main hierarchy, which is enclosed in rectangles in Fig 8, applies to models whose proteins have arbitrary amino acid compositions.^32^ Some simplifications arise when all proteins of the model have identical composition of their building blocks (Sec. II E).^31^ The hierarchy formed by such models parallels the main hierarchy, and is depicted in the right half of Fig 8.

While the assumption of uniformly composed proteins may appear to be too restricting, in fact all models that link cell metabolism to the synthesis of macromolecules through a single “translation” reaction with fixed stoichiometry (like the biomass reaction of FBA) fall into this category (e.g., refs. 27–29, 72, and 73). To this class also belong all RBA models with a single “translation capacity” constraint,^33,74^ as well as schemes that focus on the flux through a single “output” or “objective” reaction.^20,21,75^ The generalization of de Groot et al., which assigns a separate translation reaction to every enzyme,^32^ thus departs from a type of model that is deeply ingrained in the field.

In ref. 31, the optimization problem at the highest level of this parallel hierarchy was examined for models with **S** of full column rank (class 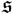 in Table 3), which assumes that only the active reactions are included. Our treatment allowed for the selection of the active reactions among all possible reactions, as illustrated by the analysis of the TVR model with two alternative pathways (Sec. II G 2 b).

The next level (highlighted in yellow in Fig 8) is especially interesting since almost all of the studies cited above are related to it either directly or indirectly. As already stated, the assumption of constant density is not needed for this optimization. In refs. 20 and 21, however, either the ribosome flux is maximized for a fixed total enzyme density,^21^ or the total enzyme density is minimized for a fixed flux through the ribosomes.^20^ Why is the macromolecular density present in these two cases, although it is absent from the optimization in the yellow rectangle of Fig 8?

In the case of models with proteins of uniform composition, both the growth time and the macromolecular density are proportional to the total protein content expressed in units of amino acids, 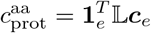 [eq. (42)].

Specifically,

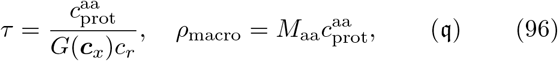

where *M*_aa_ is the molar mass of an average amino acid. Recognizing the denominator in (96) to be the flux through the ribosome, i.e., *υ*_*r*_ = *G*(***c***_*x*_)*c*_*r*_, we thus have the relationship

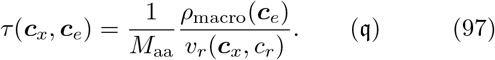

For fixed ribosome flux, minimizing the macromolecular density is seen to be equivalent to minimizing the growth time. This is the setting of ref. 20. Alternatively, if the macromolecular density is taken to be constant, maximizing *υ*_*r*_ is equivalent to minimizing the growth time, which is the setting of ref. 21.

Most significantly, for models with uniform proteins, the equality constraint of the optimization problem can be rewritten as a null-space problem [*cf.* (44)]:

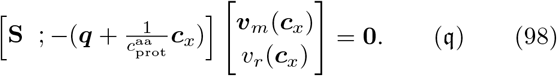

Here, [**S**; ***b***] denotes the matrix obtained by augmenting the metabolic stoichiometry matrix with an extra column equal to the vector ***b***. (This is equivalent to appending the negatives of the stoichiometric coefficients of the “biomass reaction” to the stoichiometry matrix of a metabolic model.) In (98), the column vector multiplying the augmented matrix from the right is

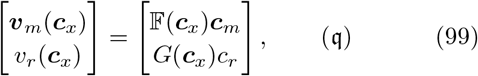

whose components are the fluxes (per unit volume) of the metabolic reactions, ***υ***_*m*_ = ***υ***^met^/*V*, and the single protein-synthesis reaction, *υ*_*r*_. Clearly, the flux vector satisfying the constraint (98) lies in the right null space of 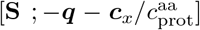. It can therefore be written as a weighted sum of the elementary flux modes (EFMs) of this matrix with non-negative weights.

For the purposes of the optimization, the augmented stoichiometric matrix in (98) is not particularly convenient since it is a function of the metabolite concentrations, which are not known ahead of time. This inconvenience is removed at Level 1 (highlighted in yellow), where the dilution of the metabolites is neglected. As the augmented matrix now simplifies to [**S**; −***q***], it can be calculated without knowledge of the metabolite concentrations, and its EFMs can be enumerated before the optimization is initiated. Relying on the results of refs. 20 and 21 that the optimal flux vector is a single EFM, and not a superposition of two or more EFMs, the optimization of the growth rate can then proceed in two steps. First, remaining on a single EFM, the metabolite concentrations are adjusted to yield the maximal growth rate. Second, the maximal growth rates of all EFMs are compared and the EFM with the highest growth rate is selected. (This approach was illustrated in ref. 21 on a simple network with only two EFMs.)

Even if the explosion of EFMs in large, genome-scale networks were not a problem, it is clear from Fig 8 that such two-stage optimization strategy will not be directly applicable when one moves up (including metabolite dilution) or to the left (allowing for proteins with different compositions) of the optimization problem in the yellow rectangle. In either case, the mass-balance equations of the metabolites cannot be written as a null-space problem of a constant matrix.

### C. Outlook

Within the framework of the presented autosynthetic cell model, *N*_*e*_ parameters were necessary to uniquely describe the states of balanced growth (Sec. II B 2). Although the choice of these parameters was not unique, the concentrations of the *N*_*e*_ enzymes were a natural option (Sec. II C 1). In the formalism of the current paper, these protein concentrations constitute the “state variables” of the cell. After either neglecting the contribution of metabolite dilution or assuming constant cell density, the number of necessary independent parameters dropped by one to *N*_*m*_. Because the growth rate of the model naturally contained the ratios between the concentrations of the metabolic enzymes and the concentration of the ribosomes [*cf.* (24)], these ratios were our variables of choice (Sec. II G).

In ref. 32, the allocation fractions of the ribosome were selected as the *N*_*m*_ independent variables. These were called the “control variables” of the cell. When the stoichiometry matrix of the cell model has a left inverse, there are *N*_*m*_ linearly independent metabolites (Sec. II C 2). The concentrations of these metabolites were selected as the independent variables in ref. 31. Since the metabolome of the cell provides a readout of its activity, we refer to the metabolite concentrations as the “readout variables” of the cell. These three choices of independent variables are illustrated schematically in Fig 9.

**Fig 9.**
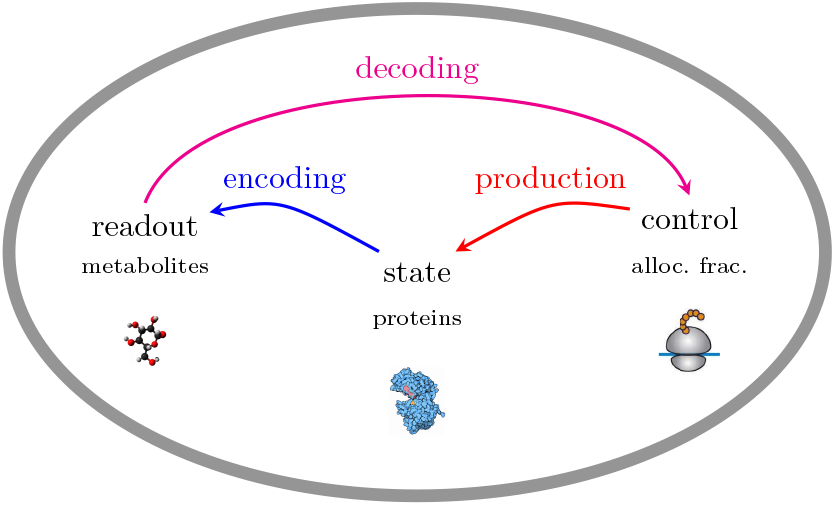
Relationships between the state variables (enzyme concentrations ***c***_*e*_), readout variables (metabolite concentrations ***c***_*x*_), and control variables (ribosome allocation fractions ***α***) of a cell.

Ultimately, the description of balanced growth should be independent of the choice of free variables. It is therefore important to elucidate the mathematical conditions that ensure this equivalence. At the same time, there could be various reasons for preferring one set of variables to the others. On the analytical side, it could be that the resulting equations are simpler and thus easier to interpret intuitively or implement numerically. On the practical side, experimental information about the three sets of variables may not be uniformly available at any specific growth condition or across different conditions. In all cases, however, even more flexibility should be gained by pooling together the strengths of the different variables and switching between them in the modeling process.

When it comes to their biological context, it is absolutely essential to consider the three alternative sets of independent variables simultaneously (Fig 9). After all, a desired protein state is generated by allocating the correct fractions of the ribosome pool to the production of the different protein types (“production” arrow in Fig 9). These allocation fractions, on the other hand, are determined with the help of proteins that regulate transcription and translation by “reading out” the metabolic status of the cell (“decoding” arrow in Fig 9). For the previous two processes to be successful, however, the cell metabolome should provide a faithful image of the enzymatic state of the cell (“encoding” arrow in Fig 9).

Such systems-level perspective on the problem enables us to ask many further questions. At the encoding stage: What are the mathematical conditions that the network of metabolic reactions should satisfy for the metabolic read-out variables to offer a faithful representation of the enzymatic state of the cell?^75^ Are there sets of several strategically chosen metabolites that can encode this state with minimal loss of information?^76^ At the decoding stage: What should the network structure of the cellular transcription/translation regulation be for the decoding from the strategically chosen metabolites to be as faithful as possible? What should the mathematical forms of the interactions between the sensed metabolites and their sensor proteins be? How well can these functional forms be approximated by actual molecular bindings? At the production stage: How is the allocation of the ribosomes to the synthesis of different proteins regulated? How does the direct involvement of metabolites (e.g., ppGpp) in this process modify the regulatory network of the decoding stage? Regarding the integration of the three stages: What mathematical properties should the resulting system have for it to respond homeostatically to different environmental perturbations?^77^ Can this system adjust to different perturbations such that, in each case, the final physiological states are good approximations to the states with optimal fitness?^29,75^

Unicellular organisms appear to have found—at least approximate—solutions to these questions through evolution. We hope that the mathematical framework of autosynthetic cell models of the type studied here and elsewhere,^31,32,78,79^ will help us gain understanding about the spectrum of all possible solutions, both actual and biologically unexplored.

## Acknowledgments

Stimulating discussions with Hugo Dourado, Ariel Amir, Xiao-Pan Hu, Tin Yau Pang, Alexander Kroll, and Ohad Golan are gratefully acknowledged. This work was funded by the Volkswagenstiftung under the “Life?” initiative, by the Deutsche Forschungsgemeinschaft (DFG, German Research Foundation) through CRC 1310 and through grant EXC 2048/1 (Project ID: 390686111) under Germany’s Excellence Strategy.

