## Supporting Information for "Structure of autosynthetic models of balanced cell growth and numerical optimization of their growth rate"

Deniz Sezer,* Peter Schubert, and Martin J. Lercher
*Institute for Computer Science and Department of Biology, Heinrich Heine University, 40221 Düsseldorf, Germany*
(Dated: September 20, 2020)

### S1. DERIVATIVES WITH RESPECT TO THE CONCENTRATIONS

In this section we provide expressions for the derivatives of the growth rate, its reciprocal, the growth time, and the constraint functions, with respect to the concentration vectors $\boldsymbol{c}_x$ (metabolites) and $\boldsymbol{c}_e$ (enzymes).

#### A. Growth rate and growth time

We start by relating the derivatives of the growth rate, $\lambda$, to the derivatives of the growth time, which was defined as

$$\tau = \lambda^{-1}. \tag{1}$$

For the derivative of the growth rate with respect to the metabolite concentrations, $\boldsymbol{c}_x$, from (1) we find that

$$\boldsymbol{\nabla}_x \lambda(\boldsymbol{c}_x, \boldsymbol{c}_e) = -\frac{1}{\tau^2}\boldsymbol{\nabla}_x \tau = -\lambda^2 \boldsymbol{\nabla}_x \tau. \tag{2}$$

In these expressions, and everywhere below, the gradient of a scalar function is to be viewed as a row vector. The row vectors $\boldsymbol{\nabla}_x \lambda$ and $\boldsymbol{\nabla}_x \tau$ in (2) have $N_x$ components, where $N_x$ denotes the number of metabolites in the model.

Similarly, the derivative of the growth rate with respect to the $N_e$ enzyme concentrations, $\boldsymbol{c}_e$, is

$$\boldsymbol{\nabla}_e \lambda(\boldsymbol{c}_x, \boldsymbol{c}_e) = -\lambda^2 \boldsymbol{\nabla}_e \tau. \tag{3}$$

The two row vectors $\boldsymbol{\nabla}_e \lambda$ and $\boldsymbol{\nabla}_e \tau$ now have $N_e$ components.

In the main text of the paper, the growth time was expressed in terms of the concentrations of the metabolites and the enzymes as

$$\tau(\boldsymbol{c}_x, \boldsymbol{c}_e) = \frac{1}{c_r}\mathbf{1}_e^T \mathbb{G}^{-1}(\boldsymbol{c}_x)\mathbb{L}\boldsymbol{c}_e. \tag{4}$$

In this expression, $c_r$ is the concentration of the ribosomes and $\mathbf{1}_e^T$ denotes a row vector all of whose $N_e$ elements are equal to one; the main diagonals of the diagonal matrices $\mathbb{G}$ and $\mathbb{L}$ contain, respectively, the ribosomal molar reaction rates for the synthesis of the $N_e$ proteins and the amino acid sequence lengths of these proteins.

When differentiating $\tau$ with respect to the concentrations of the metabolites, it is convenient to introduce the following column vector with $N_e$ components:

$$\boldsymbol{g}(\boldsymbol{c}_x) = \mathbb{L}\mathbb{G}^{-1}(\boldsymbol{c}_x)\mathbf{1}_e. \tag{5}$$

Then, the growth time in (4) becomes

$$\tau(\boldsymbol{c}_x, \boldsymbol{c}_e) = \frac{1}{c_r}\boldsymbol{g}^T(\boldsymbol{c}_x)\boldsymbol{c}_e = \frac{1}{c_r}\boldsymbol{c}_e^T \boldsymbol{g}(\boldsymbol{c}_x). \tag{6}$$

From the first equality, the derivative of the growth time with respect to the metabolite concentrations is found to be

$$\boldsymbol{\nabla}_x \tau = \frac{1}{c_r}\boldsymbol{c}_e^T[\boldsymbol{\nabla}_x \boldsymbol{g}(\boldsymbol{c}_x)], \tag{7}$$

where the components of the $N_e \times N_x$-matrix $\boldsymbol{\nabla}_x \boldsymbol{g}$ are defined as follows:

$$[\boldsymbol{\nabla}_x \boldsymbol{g}(\boldsymbol{c}_x)]_{\epsilon\xi} = \frac{\partial g_\epsilon}{\partial c_\xi}. \tag{8}$$

(The differentiation with respect to the metabolite concentration is along the second index of the matrix.)

When differentiating $\tau$ with respect to the concentrations of the enzymes, we need to consider the metabolic enzymes and the ribosomes separately. To this end, we write (6) as

$$\tau(\boldsymbol{c}_x, \boldsymbol{c}_m, c_r) = \frac{1}{c_r}\boldsymbol{g}_m^T(\boldsymbol{c}_x)\boldsymbol{c}_m + g_r(\boldsymbol{c}_x), \tag{9}$$

where $\boldsymbol{c}_m$ is a vector containing the concentrations of the $N_m$ metabolic enzymes, and $\boldsymbol{g}_m$ and $g_r$ refer to the components of the vector $\boldsymbol{g}$ that correspond to the metabolic enzymes and the ribosomes, respectively. With this separation,

$$\boldsymbol{\nabla}_m \tau = \frac{1}{c_r}\boldsymbol{g}_m^T(\boldsymbol{c}_x), \qquad \nabla_r \tau = -\frac{1}{c_r^2}\boldsymbol{g}_m^T(\boldsymbol{c}_x)\boldsymbol{c}_m, \tag{10}$$

where $\boldsymbol{\nabla}_m \tau$ is a row vector with $N_m$ components and $\nabla_r \tau$ is a scalar.

#### B. The constraint function $\boldsymbol{h}_x$

To impose the $N_x$ balanced-growth equalities of the metabolites as constraints in the optimization of the growth time, we defined the function

$$\boldsymbol{h}_x(\boldsymbol{c}_x, \boldsymbol{c}_e) = \mathbf{S}\mathbb{F}(\boldsymbol{c}_x)\boldsymbol{c}_m - \lambda(\boldsymbol{c}_x, \boldsymbol{c}_e)\left[\mathbf{P}\boldsymbol{c}_e + \boldsymbol{c}_x\right]. \tag{11}$$

Here, $\mathbf{S}$ is the stoichiometry matrix of the metabolic reactions only, $\mathbb{F}$ is a diagonal matrix containing the molar rates of the metabolic enzymes, and the $N_x \times N_e$-matrix $\mathbf{P} = \mathbf{Q}\mathbb{L}$ contains the stoichiometric coefficients of the metabolites that appear in the protein synthesis reactions.

In (11), we treat the enzyme concentrations, $\boldsymbol{c}_e$, as fixed parameters and view the metabolite concentrations, $\boldsymbol{c}_x$, as the variables on which the function depends.

To be able to write the derivative of $\boldsymbol{h}_x$ with respect to these variables in a compact form using matrix notation, it is convenient to introduce the column vector

$$\boldsymbol{f}(\boldsymbol{c}_x) = \mathbb{F}(\boldsymbol{c}_x)\mathbf{1}_m \tag{12}$$

with $N_m$ components and the $N_m \times N_x$ derivative matrix $\boldsymbol{\nabla}_x \boldsymbol{f}$, whose components are defined as

$$[\boldsymbol{\nabla}_x \boldsymbol{f}(\boldsymbol{c}_x)]_{\mu\xi} = \frac{\partial f_\mu}{\partial c_\xi}. \tag{13}$$

(Differentiation is again placed along the second index of the matrix.)

We further transform the vector $\boldsymbol{c}_m$ in (11), which is treated as constant when differentiating with respect to $\boldsymbol{c}_x$, into the diagonal matrix $\mathbb{C}_m = \mathrm{diag}(\boldsymbol{c}_m)$, and rewrite $\boldsymbol{h}_x$ as

$$\boldsymbol{h}_x(\boldsymbol{c}_x) = \mathbf{S}\mathbb{C}_m \boldsymbol{f}(\boldsymbol{c}_x) - \lambda(\boldsymbol{c}_x)[\mathbf{P}\boldsymbol{c}_e + \boldsymbol{c}_x]. \tag{14}$$

Its derivative with respect to $\boldsymbol{c}_x$ is now

$$\boldsymbol{\nabla}_x \boldsymbol{h}_x = \mathbf{S}\mathbb{C}_m[\boldsymbol{\nabla}_x \boldsymbol{f}] - [\mathbf{P}\boldsymbol{c}_e + \boldsymbol{c}_x]\boldsymbol{\nabla}_x \lambda - \lambda \mathbb{I}_x, \tag{15}$$

where $\mathbb{I}_x$ is the $N_x \times N_x$ identity matrix. If desired, the gradient of the growth rate can be replaced by the gradient of the growth time [*cf.* (2)]:

$$\boldsymbol{\nabla}_x \boldsymbol{h}_x = \mathbf{S}\mathbb{C}_m[\boldsymbol{\nabla}_x \boldsymbol{f}] + \lambda^2[\mathbf{P}\boldsymbol{c}_e + \boldsymbol{c}_x]\boldsymbol{\nabla}_x \tau - \lambda \mathbb{I}_x. \tag{16}$$

The presence of the identity matrix on the right-hand side of (15) makes it likely that, for $\lambda \neq 0$, the Jacobian matrix $\boldsymbol{\nabla}_x \boldsymbol{h}_x$ will be invertible for almost all $\boldsymbol{c}_x$ and $\boldsymbol{c}_e$, with possible exception of isolated unique combinations of these concentrations.

#### C. The constraint function $\boldsymbol{h}_m$

In the case of a stoichiometry matrix of full column rank, the $N_x$ balanced-growth equalities of the metabolites could be used to generate the $N_m$ equality constraints

$$\boldsymbol{h}_m(\boldsymbol{c}_x, \boldsymbol{c}_e) = \mathbf{0}, \tag{17}$$

where

$$\boldsymbol{h}_m(\boldsymbol{c}_x, \boldsymbol{c}_e) = \mathbb{F}(\boldsymbol{c}_x)\boldsymbol{c}_m - \lambda(\boldsymbol{c}_x, \boldsymbol{c}_e)\mathbf{S}_{\mathrm{left}}^{-1}[\mathbf{P}\boldsymbol{c}_e + \boldsymbol{c}_x], \tag{18}$$

together with the $N_{\mathrm{left}} = N_x - N_m$ conservation relations

$$\boldsymbol{h}_{\mathrm{left}}(\boldsymbol{c}_x, \boldsymbol{c}_e) = \mathbf{V}_{\mathrm{left}}[\mathbf{P}\boldsymbol{c}_e + \boldsymbol{c}_x] = \mathbf{0}. \tag{19}$$

The matrices $\mathbf{S}_{\mathrm{left}}^{-1}$ and $\mathbf{V}_{\mathrm{left}}$ in the last two equations are such that

$$\mathbf{S}_{\mathrm{left}}^{-1}\mathbf{S} = \mathbb{I}_m, \qquad \mathbf{V}_{\mathrm{left}}\mathbf{S} = 0, \tag{20}$$

where $\mathbb{I}_m$ is the $N_m \times N_m$ identity matrix.

It should be pointed out that the left inverse of the metabolic stoichiometric matrix, $\mathbf{S}$, is not unique. However, the different possible choices of $\mathbf{S}_{\mathrm{left}}^{-1}$ differ only in their action on the vectors in the left null space of $\mathbf{S}$. In our case, (19) ensures that $[\mathbf{P}\boldsymbol{c}_e + \boldsymbol{c}_x]$ is orthogonal to the left null space of $\mathbf{S}$. Since the projection of $[\mathbf{P}\boldsymbol{c}_e + \boldsymbol{c}_x]$ on the left null space vanishes, the vector $\mathbf{S}_{\mathrm{left}}^{-1}[\mathbf{P}\boldsymbol{c}_e + \boldsymbol{c}_x]$ in (18) is identical for the different choices of $\mathbf{S}_{\mathrm{left}}^{-1}$.

We consider $\boldsymbol{h}_m$ as a function of the concentrations of the metabolic enzymes, $\boldsymbol{c}_m$, and view the metabolite concentrations as parameters. From (18), the $N_m \times N_m$ derivative matrix is found to be

$$\boldsymbol{\nabla}_m \boldsymbol{h}_m = \mathbb{F} - \lambda \mathbf{S}_{\mathrm{left}}^{-1}\mathbf{P}_m + \lambda^2 \mathbf{S}_{\mathrm{left}}^{-1}[\mathbf{P}\boldsymbol{c}_e + \boldsymbol{c}_x]\boldsymbol{\nabla}_m \tau, \tag{21}$$

where the $N_x \times N_m$-matrix $\mathbf{P}_m$ is composed of the first $N_m$ columns of $\mathbf{P}$, which correspond to the metabolic enzymes.

We observe that the square matrix $\boldsymbol{\nabla}_m \boldsymbol{h}_m$ is likely to be invertible for almost all $\boldsymbol{c}_x$ and $\boldsymbol{c}_e$, as long as the diagonal matrix $\mathbb{F}$ is invertible, i.e., all its diagonal elements are non-zero. This would be the case if the substrates of all metabolic reactions are present and none of the reversible reactions is in chemical equilibrium. The former condition is met by restricting the metabolic network to the active reactions only, as was done by Dourado and Lercher.[1]

### S2. DERIVATIVES WITH RESPECT TO THE PROTEIN RATIOS

In the main text, we observed that the growth time does not depend on the overall magnitude of the concentrations of the enzymes but only on their ratios with the concentration of the ribosome, which we denoted as

$$\boldsymbol{\theta}_e = \boldsymbol{c}_e / c_r. \tag{22}$$

In this section we treat all functions from the previous section as functions of these ratios, rather than the concentrations $\boldsymbol{c}_e$. As $c_r/c_r = 1$, only the ratios of the metabolic enzymes, $\boldsymbol{\theta}_m$, are of interest.

#### A. Growth time and growth rate

In terms of the protein ratios, the growth time is given by the following dot product:

$$\tau(\boldsymbol{c}_x, \boldsymbol{\theta}_e) = \boldsymbol{g}_e^T(\boldsymbol{c}_x)\boldsymbol{\theta}_e. \tag{23}$$

Writing this growth time as [*cf.* (9)]

$$\tau(\boldsymbol{c}_x, \boldsymbol{\theta}_m) = \boldsymbol{g}_m^T(\boldsymbol{c}_x)\boldsymbol{\theta}_m + g_r(\boldsymbol{c}_x), \tag{24}$$

we find its derivatives with respect to $\boldsymbol{\theta}_m$ and $\boldsymbol{c}_x$ to be the row vectors

$$\boldsymbol{\nabla}_m\tau = \boldsymbol{g}_m^T(\boldsymbol{c}_x), \qquad \boldsymbol{\nabla}_x\tau = \boldsymbol{\theta}_e^T[\boldsymbol{\nabla}_x\boldsymbol{g}(\boldsymbol{c}_x)]. \tag{25}$$

Naturally, the derivative of the growth rate with respect to the ratios of the metabolic enzymes is

$$\boldsymbol{\nabla}_m\lambda(\boldsymbol{c}_x, \boldsymbol{\theta}_m) = -\lambda^2\boldsymbol{\nabla}_m\tau. \tag{26}$$

#### B. The constraint function $\tilde{\boldsymbol{h}}_x$

First we examine the situation with neglected metabolite dilution, where the constraint function was

$$\tilde{\boldsymbol{h}}_x(\boldsymbol{c}_x, \boldsymbol{\theta}_m) = \mathbf{S}\mathbb{F}(\boldsymbol{c}_x)\boldsymbol{\theta}_m - \lambda(\boldsymbol{c}_x, \boldsymbol{\theta}_m)\mathbf{P}\boldsymbol{\theta}_e. \tag{27}$$

The additional contribution of metabolite dilution to this function will be included in the next subsection.

The derivative of $\tilde{\boldsymbol{h}}_x$ with respect to the protein ratios, $\boldsymbol{\theta}_m$, is the $N_x \times N_m$-matrix

$$\boldsymbol{\nabla}_m\tilde{\boldsymbol{h}}_x = \mathbf{S}\mathbb{F} - \lambda\mathbf{P}_m + \lambda^2\mathbf{P}\boldsymbol{\theta}_e\boldsymbol{\nabla}_m\tau, \tag{28}$$

where (26) was used to replace the derivative of the growth rate with the derivative of the growth time in the last term.

In general, $\boldsymbol{\nabla}_m\tilde{\boldsymbol{h}}_x$ is not a square matrix. It is square only when the number of metabolites ($N_x$) is equal to the number of metabolic reactions ($N_m$). We will further assume that $\mathbf{S}$ is of full rank, hence invertible. In this special case, $\boldsymbol{\nabla}_m\tilde{\boldsymbol{h}}_x$ will be invertible when all metabolic reactions are active and out of chemical equilibrium (i.e., there are no zeros along the main diagonal of $\mathbb{F}$).

Most generally, the reduced gradient method is used with the ratios $\boldsymbol{\theta}_m$ treated as the independent variables and the concentrations $\boldsymbol{c}_x$ as the dependent variables. Then, the rectangular matrix $\boldsymbol{\nabla}_m\tilde{\boldsymbol{h}}_x$ is used as is. The square matrix $\boldsymbol{\nabla}_x\tilde{\boldsymbol{h}}_x$, on the other hand, needs to be inverted.

To derive an expression for this latter derivative matrix, we first rewrite $\tilde{\boldsymbol{h}}_x$ in (27) as

$$\tilde{\boldsymbol{h}}_x(\boldsymbol{c}_x, \boldsymbol{\theta}_m) = \mathbf{S}\Theta_m\boldsymbol{f}(\boldsymbol{c}_x) - \lambda(\boldsymbol{c}_x, \boldsymbol{\theta}_m)\mathbf{P}\boldsymbol{\theta}_e, \tag{29}$$

where the diagonal matrix $\Theta_m = \mathrm{diag}(\boldsymbol{\theta}_m)$ is treated as constant during the differentiation. The derivative with respect to the metabolite concentrations is now the $N_x \times N_x$-matrix

$$\boldsymbol{\nabla}_x\tilde{\boldsymbol{h}}_x = \mathbf{S}\Theta_m[\boldsymbol{\nabla}_x\boldsymbol{f}] + \lambda^2\mathbf{P}\boldsymbol{\theta}_e\boldsymbol{\nabla}_x\tau, \tag{30}$$

where the gradient of the growth rate was replaced by the gradient of the growth time in the last term. Since $\mathbf{P}\boldsymbol{\theta}_e$ is a column vector with $N_x$ components and $\boldsymbol{\nabla}_x\tau$ is a row vector with $N_x$ components, this last term is an $N_x \times N_x$-matrix, as it should. As the entries of the vector $\mathbf{P}\boldsymbol{\theta}_e$ are non-zero only for the metabolites that participate in the reactions of protein synthesis, this last term cannot ensure that $\boldsymbol{\nabla}_x\tilde{\boldsymbol{h}}_x$ is invertible for most combinations of $\boldsymbol{c}_x$ and $\boldsymbol{\theta}_m$; this can only be done by the first additive term on the right-hand side of (30). Thus, for $\boldsymbol{\nabla}_x\tilde{\boldsymbol{h}}_x$ to be almost always invertible, the $N_m \times N_x$-matrix $\boldsymbol{\nabla}_x\boldsymbol{f}$ should have a certain structure, correlated with the structure of the $N_x \times N_m$-matrix $\mathbf{S}$.

The metabolic stoichiometry matrix, $\mathbf{S}$, has positive entries for the products and negative entries for the substrates of the metabolic reactions (relative to the assumed forward directions). As an example, the column of $\mathbf{S}$ for a given metabolic reaction may look like this:

$$\begin{array}{c} \\ \\ c_i \\ \\ c_j \\ \\ c_k \\ c_l \\ \\ \end{array}\begin{array}{c}\mu\\ \left[\begin{array}{c} \vdots \\ -1 \\ \vdots \\ 1 \\ \vdots \\ 1 \\ -1 \\ \vdots \end{array}\right]\end{array}, \tag{31}$$

where the dots indicate entries equal to zero. Clearly, metabolites $i$ and $l$ are the substrates, while $j$ and $k$ are the products of this reaction. In general, the rate law of this reaction may look like this

$$F_\mu = \frac{k_\mu\left(\frac{c_i c_l}{K_\mu^i K_\mu^l} - \frac{c_j c_k}{K_\mu^i K_\mu^l K_\mu^{\mathrm{eq}}}\right)}{\left(1 + \frac{c_i}{K_\mu^i}\right)\left(1 + \frac{c_j}{K_\mu^j}\right)\left(1 + \frac{c_k}{K_\mu^k}\right)\left(1 + \frac{c_l}{K_\mu^l}\right)}, \tag{32}$$

where $K_\mu^{\mathrm{eq}}$ is the equilibrium constant of the reaction. Then, the corresponding row of the derivative matrix $\boldsymbol{\nabla}_x\boldsymbol{f}$ would have the following sign structure:

$$\mu\;\Big[\cdots\;\overset{c_i}{1}\;\cdots\;\overset{c_j}{-1}\;\cdots\;\overset{c_k}{-1}\;\overset{c_l}{1}\;\cdots\Big]. \tag{33}$$

Intuitively, these signs indicate that increasing the substrate concentrations would increase the reaction rate, while increasing the product concentrations would reduce the rate. Thus, up to the inversion of its overall sign, the matrix $\boldsymbol{\nabla}_x\boldsymbol{f}$ would have the same sign structure as the transpose of $\mathbf{S}$. As a result, the rank of the matrix product $\mathbf{S}\Theta_m[\boldsymbol{\nabla}_x\boldsymbol{f}]$ appearing on the right-hand side of (30) should be similar to the rank of $\mathbf{S}\mathbf{S}^T$. The latter is invertible for $\mathbf{S}$ of full row rank.

What does this imply for the invertibility of $\boldsymbol{\nabla}_x\tilde{\boldsymbol{h}}_x$?

First, there could be a problem if the rank of $\mathbf{S}$ is smaller than the number of its rows. In this case, the concentrations of some metabolites can be expressed in terms of the others through the conservation relations (19). However, as pointed out in the main text, these relations are compromised when the contribution of metabolite dilution to the balanced-growth constraints is dropped out. Thus, in the the case of dependent metabolites (i.e., $\mathbf{S}$ not of full row rank), metabolite dilution should be accounted for. The invertibility of the Jacobian matrix is then ensured by the terms that are not present in $\boldsymbol{\nabla}_x\tilde{\boldsymbol{h}}_x$ but are derived in the next subsection.

Second, even when $\mathbf{S}$ is of full row rank (i.e., the concentrations of all metabolites are linearly independent from the others), the rank of $\mathbf{S}\Theta_m[\boldsymbol{\nabla}_x\boldsymbol{f}]$ could be smaller than the rank of $\mathbf{S}\mathbf{S}^T$ if the rate laws of some reactions do not contain some of the products of these reactions.

For example, suppose that the equilibrium constant $K_\mu^{\mathrm{eq}}$ in the numerator of (32) is so large that this reaction is practically irreversible. Then, we could be tempted to approximate the irreversible rate law as

$$F_\mu = \frac{k_\mu \frac{c_i c_l}{K_\mu^i K_\mu^l}}{\left(1+\frac{c_i}{K_\mu^i}\right)\left(1+\frac{c_l}{K_\mu^l}\right)}. \qquad (34)$$

If so, the corresponding row of the derivative matrix $\boldsymbol{\nabla}_x\boldsymbol{f}$ would have the following sign structure:

$$\mu\ \left[\begin{array}{ccccccccc} \cdots & \overset{c_i}{1} & \cdots & \overset{c_j}{0} & \cdots & \overset{c_k}{0} & \overset{c_l}{1} & \cdots \end{array}\right]. \qquad (35)$$

Naturally, we now have zeros for the *products* of the reaction since these do not appear in the rate law. If the metabolites $j$ and $k$ also do not appear in any other reaction of the network, then the $j$-th and $k$-th diagonal entries of $\mathbf{S}\Theta_m[\boldsymbol{\nabla}_x\boldsymbol{f}]$ would be equal to zero. This is likely to reduce the rank of the matrix, compared to the rank of $\mathbf{S}\mathbf{S}^T$, and thus compromise the invertibility of the Jacobian. (As an example of a metabolite that is produced but not used by any other reaction, we could think of acetate production during fermentation in, for example, models of overflow metabolism. Since acetate is not exported by a dedicated transporter proteins but diffuses freely through the lipid component of the cell membrane, an explicit reaction that takes acetate as a substrate may be missing from the active metabolic network.)

Compared to the rate law (32), the *products* of the reaction are missing from both the numerator and the denominator of (34). While removing the product concentrations from the numerator is part of the assumption that the reaction is irreversible, removing them from the denominator is actually not required by the irreversibility. In fact, we could have written the rate law of the irreversible reaction as

$$F_\mu = \frac{k_\mu \frac{c_i c_l}{K_\mu^i K_\mu^l}}{\left(1+\frac{c_i}{K_\mu^i}\right)\left(1+\frac{c_j}{K_\mu^j}\right)\left(1+\frac{c_k}{K_\mu^k}\right)\left(1+\frac{c_l}{K_\mu^l}\right)}. \qquad (36)$$

In this case, the sign structure of the derivative would be the same as that shown in (33), thus leading to $\mathbf{S}\Theta_m[\boldsymbol{\nabla}_x\boldsymbol{f}]$ that is of full rank for almost all combinations of the variables $\boldsymbol{c}_x$ and $\boldsymbol{\theta}_m$. By retaining the factors in the denominator of an irreversible rate law, we account for the inhibition of the enzyme by the products of the reaction. The general importance of *product inhibition* is extensively discussed in Cornish-Bowden's excellent textbook on enzyme kinetics.[2] Its role as a feedback that stabilizes the steady state of a metabolic network against metabolite perturbation is discussed in ref. 3 (Sec. II 5).

### S3. INCLUDING METABOLITE DILUTION WITH A DENSITY CONSTRAINT

#### A. The constraint function

When the contribution of metabolite dilution is included, the constraint function $\tilde{\boldsymbol{h}}_x$ acquires the extra additive term

$$-\frac{\lambda}{c_r}\boldsymbol{c}_x. \qquad (37)$$

As shown in the main text, the concentration of the ribosome appearing in the denominator of (37) may be expressed in terms of the optimization variables $\boldsymbol{c}_x$ and $\boldsymbol{\theta}_m$ if either the macromolecular or the dry-mass density of the cell is assumed to be constant. These two cases are examined next.

In the case of constant macromolecular density, $\rho_{\mathrm{macro}}$, the constraint function $\tilde{\boldsymbol{h}}_x$ is modified as

$$\begin{aligned}\tilde{\boldsymbol{h}}_x^{\mathrm{macro}}(\boldsymbol{c}_x,\boldsymbol{\theta}_m) &= \tilde{\boldsymbol{h}}_x(\boldsymbol{c}_x,\boldsymbol{\theta}_m)\\ &\quad - \lambda(\boldsymbol{c}_x,\boldsymbol{\theta}_m)\rho_{\mathrm{macro}}^{-1}\boldsymbol{\mu}_e^T\boldsymbol{\theta}_e\boldsymbol{c}_x,\end{aligned} \qquad (38)$$

where we have introduced the column vector

$$\boldsymbol{\mu}_e = \mathbb{M}_e\mathbf{1}_e \qquad (39)$$

containing the molar masses of the proteins. [Recall that $\mathbb{M}_e$ was a diagonal matrix composed of the protein molar masses, and note that the dot product $\boldsymbol{\mu}_e^T\boldsymbol{\theta}_e$ in (38) produces a scalar.]

For assumed constant dry-mass density, $\rho_{\mathrm{dry}}$, we have

$$\begin{aligned}\tilde{\boldsymbol{h}}_x^{\mathrm{dry}}(\boldsymbol{c}_x,\boldsymbol{\theta}_m) &= \tilde{\boldsymbol{h}}_x(\boldsymbol{c}_x,\boldsymbol{\theta}_m)\\ &\quad - \lambda(\boldsymbol{c}_x,\boldsymbol{\theta}_m)\frac{\rho_{\mathrm{dry}}^{-1}\boldsymbol{\mu}_e^T\boldsymbol{\theta}_e}{1-\rho_{\mathrm{dry}}^{-1}\boldsymbol{\mu}_x^T\boldsymbol{c}_x}\boldsymbol{c}_x,\end{aligned} \qquad (40)$$

where we have introduced the column vector of metabolic molar masses

$$\boldsymbol{\mu}_x = \mathbb{M}_x\mathbf{1}_x. \qquad (41)$$

These two cases can be treated together after introducing the scalar function

$$\sigma(\boldsymbol{c}_x) = \begin{cases} \rho_{\mathrm{macro}}^{-1}, & \rho_{\mathrm{macro}} = \text{const.}\\ \dfrac{\rho_{\mathrm{dry}}^{-1}}{1-\rho_{\mathrm{dry}}^{-1}\boldsymbol{\mu}_x^T\boldsymbol{c}_x}, & \rho_{\mathrm{dry}} = \text{const.}\end{cases} \qquad (42)$$

and writing the vector-valued constraint functions (38) and (40) collectively as

$$\tilde{\boldsymbol{h}}_x^{\rho}(\boldsymbol{c}_x, \boldsymbol{\theta}_m) = \tilde{\boldsymbol{h}}_x(\boldsymbol{c}_x, \boldsymbol{\theta}_m) - \lambda(\boldsymbol{c}_x, \boldsymbol{\theta}_m)\sigma(\boldsymbol{c}_x)\boldsymbol{\mu}_e^T \boldsymbol{\theta}_e \boldsymbol{c}_x. \tag{43}$$

For the multiplicative update of the metabolite concentrations, $\tilde{\boldsymbol{h}}_x^{\rho}$ is separated into positive and negative contributions as follows:

$$\begin{aligned}\tilde{\boldsymbol{h}}_x^{\rho+} &= (\mathbf{S}^+\mathbb{F}^+ + \mathbf{S}^-\mathbb{F}^-)\boldsymbol{\theta}_m + \lambda\mathbf{P}^-\boldsymbol{\theta}_e \\ \tilde{\boldsymbol{h}}_x^{\rho-} &= (\mathbf{S}^+\mathbb{F}^- + \mathbf{S}^-\mathbb{F}^+)\boldsymbol{\theta}_m + \lambda\mathbf{P}^+\boldsymbol{\theta}_e + \lambda\sigma\boldsymbol{\mu}_e^T\boldsymbol{\theta}_e\boldsymbol{c}_x.\end{aligned} \tag{44}$$

Only the very last term on the right-hand side of the second line is new compared to the case with neglected metabolite dilution, which was analyzed in the main text.

#### B. Derivatives of the constraint

Because $\sigma$ is not a function of $\boldsymbol{\theta}_m$ for both density constraints, the derivative of $\tilde{\boldsymbol{h}}_x^{\rho}$ with respect to the protein ratios is the $N_x \times N_m$-matrix

$$\boldsymbol{\nabla}_m\tilde{\boldsymbol{h}}_x^{\rho} = \boldsymbol{\nabla}_m\tilde{\boldsymbol{h}}_x + \lambda\sigma\boldsymbol{c}_x[\lambda\boldsymbol{\mu}_e^T\boldsymbol{\theta}_e\boldsymbol{\nabla}_m\tau - \boldsymbol{\mu}_m^T], \tag{45}$$

with $\boldsymbol{\nabla}_m\tilde{\boldsymbol{h}}_x$ as given in (28). (This matrix need not be inverted in the reduced gradient method with $\boldsymbol{\theta}_m$ as independent variables.) Note that for $\rho_{\text{macro}}^{-1} = 0$ or $\rho_{\text{dry}}^{-1} = 0$, the derivative in (45) reduces to the case of neglected metabolite dilution.

The derivative of $\tilde{\boldsymbol{h}}_x^{\rho}$ with respect to the metabolite concentrations is the $N_x \times N_x$-matrix

$$\begin{aligned}\boldsymbol{\nabla}_x\tilde{\boldsymbol{h}}_x^{\rho} = \boldsymbol{\nabla}_x\tilde{\boldsymbol{h}}_x &- \lambda\sigma\boldsymbol{\mu}_e^T\boldsymbol{\theta}_e\mathbb{I}_x \\ &+ \lambda^2\sigma\boldsymbol{\mu}_e^T\boldsymbol{\theta}_e\boldsymbol{c}_x\boldsymbol{\nabla}_x\tau - \lambda\boldsymbol{\mu}_e^T\boldsymbol{\theta}_e\boldsymbol{c}_x\boldsymbol{\nabla}_x\sigma,\end{aligned} \tag{46}$$

with $\boldsymbol{\nabla}_x\tilde{\boldsymbol{h}}_x$ as given above in (30). Since $\boldsymbol{\nabla}_x\sigma = 0$ for constant macromolecular density, we immediately find

$$\boldsymbol{\nabla}_x\tilde{\boldsymbol{h}}_x^{\text{macro}} = \boldsymbol{\nabla}_x\tilde{\boldsymbol{h}}_x + \lambda\rho_{\text{macro}}^{-1}\boldsymbol{\mu}_e^T\boldsymbol{\theta}_e[\lambda\boldsymbol{c}_x\boldsymbol{\nabla}_x\tau - \mathbb{I}_x]. \tag{47}$$

In the case of constant dry-mass density, on the other hand, we get

$$\boldsymbol{\nabla}_x\tilde{\boldsymbol{h}}_x^{\text{dry}} = \boldsymbol{\nabla}_x\tilde{\boldsymbol{h}}_x + \lambda\sigma\boldsymbol{\mu}_e^T\boldsymbol{\theta}_e[\lambda\boldsymbol{c}_x\boldsymbol{\nabla}_x\tau - \mathbb{I}_x - \sigma\boldsymbol{c}_x\boldsymbol{\mu}_x^T], \tag{48}$$

with $\sigma$ as given in the second line of (42). Again, for $\rho_{\text{macro}}^{-1} = 0$ or $\rho_{\text{dry}}^{-1} = 0$ the derivatives in (47) and (48) reduce to the expressions for neglected metabolite dilution.

The identity matrix $\mathbb{I}_x$ that appears in both (47) and (48), ensures that $\boldsymbol{\nabla}_x\tilde{\boldsymbol{h}}_x^{\rho}$ will be invertible for almost all combinations of $\boldsymbol{c}_x$ and $\boldsymbol{\theta}_m$, even in the case of linearly dependent metabolites that are constrained by conservation relations.

### S4. FURTHER ILLUSTRATIVE EXAMPLES

#### A. Linearly dependent metabolites

The simple models TR, TVR and TUR, which were examined numerically in the main text, did not illustrate the case of having more metabolites than the rank of the stoichiometric matrix. In this case, the conservation relations that relate the metabolite concentrations to each other and to the concentrations of the proteins are compromised when the dilution of the metabolites is neglected, as we saw above.

Here, we illustrate this situation using the simple model given in the first row of fig. S1, which could be viewed as a hybrid between the TR model with a single transporter and the TUR model with two amino acids. In this model, which we call T2R for brevity, the two amino acids are internalized at a fixed ratio by the transporter. The metabolic stoichiometric matrix of the model is

$$\mathbf{S} = \begin{bmatrix} s_{11} \\ s_{21} \end{bmatrix}, \tag{49}$$

with $s_{11} > 0$ and $s_{21} > 0$. For our purposes, what matters is that the matrix $\mathbf{S}$ has a left null space. Up to an arbitrary normalization, the vectors in this one-dimensional null space look like this:

$$\boldsymbol{v}_{\text{left}}^T = \begin{bmatrix} s_{21}, & -s_{11} \end{bmatrix}. \tag{50}$$

As in the TUR model of the main text, we will take the fractional amino acid composition of the proteins to be

$$\mathbf{Q} = \begin{array}{c} \\ A \\ B \end{array}\!\!\begin{array}{c} \begin{array}{cc} T & R \end{array} \\ \left[\begin{array}{cc} 2/3 & 1/4 \\ 1/3 & 3/4 \end{array}\right] \end{array}. \tag{51}$$

Like in the TUR model, we will assume that the lengths of the two proteins are $\ell_T = 60 \times 300$ and $\ell_R = 60 \times 400$. Thus,

$$\mathbf{P} = \mathbf{Q}\mathbb{L} = \begin{array}{c} \\ A \\ B \end{array}\!\!\begin{array}{c} \begin{array}{cc} T & R \end{array} \\ \left[\begin{array}{cc} 12000 & 6000 \\ 6000 & 18000 \end{array}\right] \end{array}. \tag{52}$$

For simplicity, we will take the molar masses of the two metabolites to be equal to each other, i.e., $M_B = M_A$.

The constraint function for the T2R model ensuring the mass-balance of the metabolites is

$$\boldsymbol{h}_x = \mathbf{S}F_Tc_T - \lambda\left(\mathbf{P}\begin{bmatrix} c_T \\ c_R \end{bmatrix} + \begin{bmatrix} c_A \\ c_B \end{bmatrix}\right). \tag{53}$$

Due to the presence of a left null space of $\mathbf{S}$, the concentrations of the two metabolites should satisfy the conservation relation

$$\boldsymbol{v}_{\text{left}}^T\left(\mathbf{P}\begin{bmatrix} c_T \\ c_R \end{bmatrix} + \begin{bmatrix} c_A \\ c_B \end{bmatrix}\right) = 0. \tag{54}$$

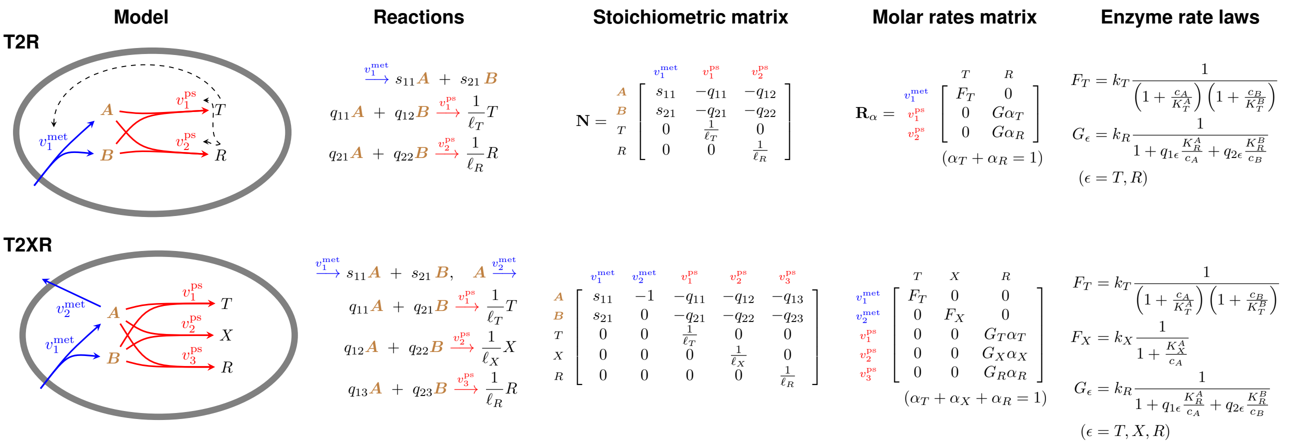


FIG. S1. Minimal autosynthetic cell models with linearly dependent metabolites (T2R) and metabolite excretion (T2XR). The metabolic reactions are colored blue, the protein-synthesis reactions are red. Product occupancy is included in the rate law of the transporter $T$, but not of the ribosome $R$ nor the exporter $X$. The rates $k_T = 100\,\mathrm{s}$, $k_R = 20\,\mathrm{s}$, $k_X = 80\,\mathrm{s}$ and Michaelis constants $K_T^A = 100\,\mathrm{mM}$, $K_T^B = 70\,\mathrm{mM}$, $K_R^A = 10\,\mathrm{mM}$, $K_R^B = 5\,\mathrm{mM}$, $K_X^A = 30\,\mathrm{mM}$ were used in the numerical examples reported in Tables S1 and S2.

We note that the first vector in the parenthesis, namely

$$\mathbf{P}\begin{bmatrix} c_T \\ c_R \end{bmatrix} = \begin{bmatrix} c_A^{\text{prot}} \\ c_B^{\text{prot}} \end{bmatrix}, \tag{55}$$

contains the concentrations of the metabolites that have already been incorporated into the proteins. The conservation relation is thus

$$\boldsymbol{v}_{\text{left}}^T \begin{bmatrix} c_A + c_A^{\text{prot}} \\ c_B + c_B^{\text{prot}} \end{bmatrix} = 0. \tag{56}$$

For concreteness, let us consider the model with

$$\mathbf{S} = \begin{bmatrix} 1 \\ 2 \end{bmatrix}, \tag{57}$$

which corresponds to the transport of two molecules of $B$ for every molecule of $A$. The conservation relation in this case,

$$2(c_A + c_A^{\text{prot}}) = c_B + c_B^{\text{prot}}, \tag{58}$$

states that the *total* concentration of $B$ inside the cell, including both free $B$ and $B$ that is already part of the proteins, should be twice as large as the *total* concentration of $A$ inside the cell. Given the stoichiometry with which $A$ and $B$ enter the cell through the transporter, this requirement makes perfect sense.

Ignoring the metabolite dilution amounts to dropping the concentrations of the metabolites from the constraint $\boldsymbol{h}_x$ in (53). When this term is neglected, the conservation relation turns into the requirement

$$\boldsymbol{v}_{\text{left}}^T \mathbf{P} \begin{bmatrix} c_T \\ c_R \end{bmatrix} = 0, \tag{59}$$

which completely fails to relate the concentrations of the metabolites to each other, but instead imposes a relationship between the concentrations of the two proteins. This example should clarify why, in practice, metabolite dilution cannot be dropped when the number of rows in the metabolic stoichiometric matrix is larger than its rank.

To retain metabolite dilution, we will assume that either the dry mass density or the macromolecular density is constant. After dividing the function $\boldsymbol{h}_x$ by the concentration of the ribosomes and expressing this concentration in terms of the other variables, we obtain the new constraint function [*cf.* (53)]

$$\tilde{\boldsymbol{h}}^\rho = \mathbf{S} F_T \theta_T - \lambda \left\{ \mathbf{P} \begin{bmatrix} \theta_T \\ 1 \end{bmatrix} + \sigma \left[ M_T, \;\; M_R \right] \begin{bmatrix} \theta_T \\ 1 \end{bmatrix} \begin{bmatrix} c_A \\ c_B \end{bmatrix} \right\}. \tag{60}$$

The separation of this function into non-negative components according to (44) implies the following multiplicative update of the metabolite concentrations:

$$\begin{bmatrix} c_A \\ c_B \end{bmatrix} \leftarrow \begin{bmatrix} c_A \\ c_B \end{bmatrix} \circ \left( \frac{\tau\, \mathbf{S} F_T \theta_T}{\mathbf{P} \begin{bmatrix} \theta_T \\ 1 \end{bmatrix} + \sigma \left[ M_T, \;\; M_R \right] \begin{bmatrix} \theta_T \\ 1 \end{bmatrix} \begin{bmatrix} c_A \\ c_B \end{bmatrix}} \right)^{\kappa} \tag{61}$$

and $\kappa > 0$. Recall that

$$\sigma = \rho_{\text{macro}}^{-1} \tag{62}$$

in the case of constant macromolecular density, and

$$\sigma = \frac{\rho_{\text{dry}}^{-1}}{1 - \rho_{\text{dry}}^{-1}(M_A c_A + M_B c_B)} \tag{63}$$

in the case of constant dry mass density.

We used the reduced gradient method, with the ratio $\theta_T$ as the independent variable, to optimize T2R models with different stoichiometries of the transporter. The results are compiled in Table S1. The upper half of the

TABLE S1. Optima of the T2R model for different stoichiometries of the transporter ($s_{11} : s_{21}$) with either constant macromolecular density ($\rho_{\mathrm{macro}} = 300\,\mathrm{g\,l^{-1}}$) or constant dry mass density ($\rho_{\mathrm{dry}} = 300\,\mathrm{g\,l^{-1}}$). (Largest $\kappa$ still converging for maximum of 800 iteration steps.)

| | | $\lambda/\mathrm{h}^{-1}$ | $\theta_T$ | $c_A/\mathrm{mM}$ | $c_B/\mathrm{mM}$ | $c_R/\mu\mathrm{M}$ | $\kappa$ |
|---|---|---|---|---|---|---|---|
| macro | 1:3 | 0.8253 | 0.1349 | 1.257 | 463.2 | 113.52 | 1.1 |
| | 1:2 | 2.3252 | 0.2464 | 110.8 | 56.59 | 105.50 | 1.4 |
| | 1:1 | 1.6036 | 0.9287 | 504.3 | 30.68 | 73.68 | 0.8 |
| | 2:1 | 1.0837 | 1.9056 | 1581. | 18.50 | 51.46 | 0.8 |
| dry | 1:3 | 1.0020 | 0.5718 | 3.434 | 1006. | 58.04 | 1.0 |
| | 1:2 | 2.3273 | 0.2434 | 108.9 | 56.06 | 99.89 | 1.4 |
| | 1:1 | 1.6434 | 0.8637 | 462.1 | 29.63 | 63.43 | 0.6 |
| | 2:1 | 1.2422 | 1.4714 | 1134. | 18.46 | 36.59 | 0.2 |

table is for constant $\rho_{\mathrm{macro}}$ and the lower half for constant $\rho_{\mathrm{dry}}$. In either case, for stoichiometry of 1:3, the concentration of $B$ at the optimum is way higher than that of $A$. Similarly, for a stoichiometry of 2:1, the concentration of $A$ at the optimum is much higher than that of $B$. Since a lot of unused precursor is accumulated in the cell, the optimal growth rates for these unbalanced stoichiometries are relatively low.

The best growth rates are obtained for a transporter stoichiometry of 1:2. The reason lies in the specific choice of the protein composition matrix $\mathbf{Q}$ in (51) and the kinetic rates of the two enzymes. Since the ribosome is slower, the cell needs more ribosomes than transporters, as evidenced by the optimal ratio $\theta_T$ for 1:2. At the same time, the ribosome is 75% B and 25% A, while the transporter is 67% A and 33% B. As a result, the total demand of the proteome for $B$ is larger than that for $A$.

The last column of Table S1 shows the largest value of the parameter $\kappa$, appearing in the multiplicative update of the dependent metabolite variables (61), for which the numerical values of the metabolite concentrations converged to within $10^{-8}$ in less than 800 iteration steps. ($\kappa$ was kept constant throughout the optimization. The reported values were determined manually.)

The first observation we make is that larger $\kappa$, hence faster "learning rate", can be used when the $A$:$B$ stoichiometry of the transporter is best matched by the internal demand of the proteome for $A$ and $B$. Deviations from this perfect match require smaller values of $\kappa$ for convergence. The second point that catches attention is that, in the more challenging cases of 1:1 and 2:1, $\kappa$ remains reasonably close to 1 in the case of constant macromolecular density, but needs to be reduced substantially in the case of constant dry-mass density. The origin of the poorer convergence in these latter cases lies in the form of $\sigma$ in (63). While $\sigma$, and hence the denominator of (61), is guaranteed to be positive when $\sigma = \rho_{\mathrm{macro}}$, it could become negative during the multiplicative updates in the case of constant dry mass density if the product $\rho_{\mathrm{dry}}^{-1}(M_A c_A + M_B c_B)$ exceeds 1. This problem was indeed encountered when larger values of $\kappa$ were used during the optimizations in the last two rows of Table S1. For this reason, the proposed multiplicative update is expected to have better convergence properties when used with a constraint on the macromolecular density, as opposed to the dry mass density, especially when the concentrations of some metabolites become unreasonably large.

#### B. Metabolite excretion

For all minimal models examined numerically in the paper, including the T2R model above, the entries of the metabolic stoichiometric matrix were non-negative. As a result, the matrix $\mathbf{S}^-$ in the non-negative decomposition

$$\mathbf{S} = \mathbf{S}^+ - \mathbf{S}^- \tag{64}$$

was never present. Since, in addition, all our reactions were irreversible, the matrix $\mathbb{F}^-$ in the non-negative decomposition

$$\mathbb{F} = \mathbb{F}^+ - \mathbb{F}^- \tag{65}$$

was also not present. Hence, the contribution $(\mathbf{S}^+\mathbb{F}^- + \mathbf{S}^-\mathbb{F}^+)\boldsymbol{\theta}_m$ to the function $\tilde{\boldsymbol{h}}_x^{\rho-}$, which appears in the denominator of the multiplicative update (44), was never active.

Here, we modify the T2R model such that it now includes an extra transporter that can expel the metabolite $A$ from the cell. The resulting T2XR model was shown in the second row of fig. S1. Its metabolic stoichiometric matrix contains a negative entry:

$$\mathbf{S} = \begin{bmatrix} s_{11} & -1 \\ s_{21} & 0 \end{bmatrix}. \tag{66}$$

Since $\mathbf{S}$ is now of full column rank ($s_{11} > 0$ and $s_{21} > 0$), there are no conservation relations between the concentrations of the two metabolites. Thus, in principle, it should also be possible to completely neglect the dilution of the metabolites in the optimization of the growth rate.

For the numerical analysis, we chose the amino acid compositions and lengths of the proteins $T$, $X$ and $R$ to be identical to those of the TUR model in the main text, with $X$ replacing $U$. Optimizations were carried out for the same four $A$:$B$ stoichiometries as before. The results for neglected metabolite dilution (i.e., no density constraint), and constraints on either $\rho_{\mathrm{macro}}$ or $\rho_{\mathrm{dry}}$ are collected in Table S2.

For the stoichiometries where $A$ accumulated in the cell before (2:1 and 1:1 in Table S1), it now pays off to allocate a fraction of the proteome to the exporter $X$ ($\theta_X$ in Table S2). This substantially reduces the internal concentrations of $A$ compared to those of the T2R model without an exporter. As a result, even the stoichiometry 2:1 with constant dry mass density converged for $\kappa$ as large as 1.6. In these two cases (i.e., 2:1 and 1:1),

TABLE S2. Same as Table S1 but for the T2XR model.

| | | $\lambda$/h$^{-1}$ | $\theta_T$ | $\theta_X$ | $c_A$/mM | $c_B$/mM | $c_R/\mu$M |
|---|---|---|---|---|---|---|---|
| none | 1:3 | x | x | x | x | x | - |
| | 1:2 | x | x | x | x | x | - |
| | 1:1 | 2.0015 | 0.3142 | 0.1056 | 65.65 | 42.90 | - |
| | 2:1 | 1.7784 | 0.3492 | 0.2822 | 89.55 | 41.19 | - |
| macro | 1:3 | 0.8253 | 0.1349 | 1e-100 | 1.257 | 463.2 | 113.52 |
| | 1:2 | 2.3465 | 0.1938 | 0.0099 | 68.71 | 54.70 | 108.32 |
| | 1:1 | 1.9990 | 0.3164 | 0.1010 | 68.19 | 40.12 | 94.29 |
| | 2:1 | 1.7748 | 0.3509 | 0.2761 | 93.57 | 37.82 | 83.07 |
| dry | 1:3 | 1.0020 | 0.5718 | 1e-272 | 3.434 | 1006. | 58.04 |
| | 1:2 | 2.3466 | 0.1940 | 0.0096 | 68.90 | 54.54 | 103.87 |
| | 1:1 | 1.9989 | 0.3165 | 0.1008 | 68.27 | 40.03 | 90.89 |
| | 2:1 | 1.7747 | 0.3509 | 0.2758 | 93.71 | 37.68 | 79.44 |

the optimal growth rates, the protein ratios, and even the metabolite concentrations are very similar for the three examined possibilities of neglected metabolite dilution ('none'), constant macromolecular density ('macro'), and constant dry mass density ('dry') in Table S2.

In the more balanced case of transporter stoichiometry 1:2, the necessary concentration of the exporter $X$ is only 1% of the concentration of the ribosome ($\theta_X$ in Table S2). Nevertheless, this is sufficient to reduce the intracellular concentrations of $A$ by about 30% compared to those in Table S1.

When the balance between $A$ and $B$ is disturbed in favor of $B$ (stoichiometry 1:3), it no longer pays off to synthesize the exporter of $A$. This is nicely predicted by the numerical optimizations with constraint on the density ('macro' and 'dry' in Table S2), in which the numerical value of $\theta_X$ was suppressed to less than $10^{-100}$!

Because the cases of vanishingly small ratio $\theta_X$ degenerate to the T2R model of the previous subsection, the optimization with neglected metabolite dilution ('none') failed to converge for the stoichiometries 1:2 and 1:3. As discussed before, dropping the dilution of the metabolites in this case impairs the existing linear relation between the metabolite concentrations.

The two simple examples in this section demonstrate that the convergence properties of the proposed multiplicative update are closely tied to how realistic the model is. Numerical difficulties are bound to be encountered when the concentrations of some metabolites become unreasonably large. Mechanisms that may naturally restrict the metabolite concentrations, like product inhibition, utilization by other enzymes, or excretion, should be included in the model. Relying on the constraint $\rho_{\text{dry}} = \text{const.}$ for suppressing the concentrations of the metabolites may compromise the convergence due to the possibility that the denominator of $\sigma$ in (42) becomes negative.

### S5. RETAINING THE RIBOSOME ALLOCATION FRACTIONS

As discussed in the main text, the ribosome allocation fractions, $\boldsymbol{\alpha}$, were retained in the analysis of ref. 4, whereas the concentrations of the metabolic enzymes were eliminated. Here, we pursue this choice and illustrate that it leads to a more complex dependence of the constraint function on the metabolite concentrations, which makes the differentiation more involved.

The balanced-growth equalities of the proteins were

$$\lambda \boldsymbol{c}_e = \mathbb{L}^{-1}\mathbb{G}(\boldsymbol{c}_x)\boldsymbol{\alpha} c_r. \tag{67}$$

The last of these $N_m + 1$ scalar equalities provides an expression for the growth rate in terms of the concentrations of the metabolites and the fraction of ribosomes working on the synthesis of new ribosomes, $\alpha_r$:

$$\lambda = \ell_r^{-1} G_r(\boldsymbol{c}_x)\alpha_r. \tag{68}$$

In ref. 4, the equalities (67) were used to eliminate the concentrations of the metabolic enzymes from the optimization problem by expressing them in terms of the other optimization variables. For the ratios between these concentrations and the concentration of the ribosome, the result is

$$\boldsymbol{\theta}_e(\boldsymbol{c}_x, \boldsymbol{\alpha}) = \frac{1}{\lambda}\mathbb{L}^{-1}\mathbb{G}(\boldsymbol{c}_x)\boldsymbol{\alpha}. \tag{69}$$

Since $\theta_r = 1$, the last element of this vector equality simply recovers the growth rate in (68). The other $N_m$ equalities can be written for the concentrations of the metabolic enzymes as

$$\boldsymbol{c}_m(\boldsymbol{c}_x, c_r, \boldsymbol{\alpha}) = c_r \frac{\mathbb{L}_m^{-1}\mathbb{G}_m(\boldsymbol{c}_x)\boldsymbol{\alpha}_m}{\lambda}, \tag{70}$$

where $\mathbb{L}_m$, $\mathbb{G}_m$ and $\boldsymbol{\alpha}_m$ refer to the components of $\mathbb{L}$, $\mathbb{G}$ and $\boldsymbol{\alpha}$ that correspond to the metabolic enzymes only.

The balanced-growth equalities of the metabolites were

$$\lambda \boldsymbol{c}_x = \mathbf{S}\mathbb{F}(\boldsymbol{c}_x)\boldsymbol{c}_m - \lambda \mathbf{Q}\mathbb{G}(\boldsymbol{c}_x)\boldsymbol{\alpha} c_r. \tag{71}$$

Substituting $\boldsymbol{c}_m$ from (70), these become

$$\lambda \boldsymbol{c}_x = \frac{c_r}{\lambda}\mathbf{S}\mathbb{F}(\boldsymbol{c}_x)\mathbb{L}_m^{-1}\mathbb{G}_m(\boldsymbol{c}_x)\boldsymbol{\alpha}_m - \lambda c_r \mathbf{Q}\mathbb{G}(\boldsymbol{c}_x)\boldsymbol{\alpha}. \tag{72}$$

Dividing both sides by $c_r$ and multiplying by $\lambda$, the corresponding constraint function can be rewritten as

$$\boldsymbol{h}_x(\boldsymbol{c}_x; \boldsymbol{\alpha}) = \mathbf{S}\mathbb{F}(\boldsymbol{c}_x)\mathbb{L}_m^{-1}\mathbb{G}_m(\boldsymbol{c}_x)\boldsymbol{\alpha}_m - \lambda^2\left[\frac{\boldsymbol{c}_x}{c_r} + \mathbf{Q}\mathbb{G}(\boldsymbol{c}_x)\boldsymbol{\alpha}\right]. \tag{73}$$

Its derivative with respect to the metabolite concentrations, $\boldsymbol{c}_x$, is more complicated than the expressions given above.

In the case of constrained dry mass density or macromolecular density, the ribosome concentration could be expressed in terms of the other variables as follows:

$$c_r^{-1}(\boldsymbol{c}_x, \boldsymbol{\theta}_e) = \sigma(\boldsymbol{c}_x)\boldsymbol{\mu}_e^T\boldsymbol{\theta}_e, \tag{74}$$

where the function $\sigma$ was defined in (42). Switching to the ribosome allocation fractions using (69), we get

$$c_r^{-1}(\boldsymbol{c}_x, \boldsymbol{\alpha}) = \frac{1}{\lambda}\sigma(\boldsymbol{c}_x)\boldsymbol{\mu}_e^T \mathbb{L}^{-1}\mathbb{G}(\boldsymbol{c}_x)\boldsymbol{\alpha}. \qquad (75)$$

Again, differentiating this expression with respect to $\boldsymbol{c}_x$ is more laborious than differentiating (74).

*

[1] H. Dourado and M. J. Lercher, Nature Communications **11**, 1226 (2020).
[2] A. Cornish-Bowden, *Fundamentals of Enzyme Kinetics*, 4th ed. (Wiley-Blackwell, 2012).
[3] J. G. Reich and E. E. Selkov, *Energy Metabolism of the Cell: A Theoretical Treatise* (Academic Press, 1981).
[4] D. H. de Groot, J. Hulshof, B. Teusink, F. J. Bruggeman, and R. Planqué, PLOS Computational Biology **16**, e1007559 (2020).
